# HSCs and Tregs cooperate to preserve extramedullary hematopoiesis under chronic inflammation

**DOI:** 10.1101/2025.02.05.636492

**Authors:** Maria Kuzmina, Srdjan Grusanovic, Jiri Brezina, Mirko Milosevic, Karolina Vanickova, Nataliia Pavliuchenko, Sarka Ruzickova, Jakub Rohlena, Dominik Filipp, Katerina Rohlenova, Tomas Brdicka, Meritxell Alberich-Jorda

## Abstract

Hematopoietic stem cells (HSCs) are localized within specialized niches of the bone marrow (BM). However, during hematological disorders or infections, the functionality of HSCs in the BM is compromised, leading to extramedullary hematopoiesis (EMH). Chronic inflammation drives EMH, yet its impact on HSCs outside the BM is poorly understood. Using a mouse model of chronic autoinflammatory disease, we demonstrated the presence of extramedullary HSCs in blood, spleen, and inflamed tails and paws. Single-cell transcriptomics revealed a unique expression profile in extramedullary HSCs, with significant upregulation of *Cd53*, MHCII-associated, and immunosuppressive genes. We further demonstrated that extramedullary CD53+ HSCs act as antigen-presenting cells, promoting the development of regulatory T cells (Tregs) to control chronic inflammation at extramedullary sites. Conversely, Tregs exert a protective role on extramedullary HSCs. Altogether, our findings revealed a mutually supportive relationship between a unique subset of HSCs and T cells in inflamed tissues during chronic inflammation.

## INTRODUCTION

Hematopoietic stem cells (HSCs) are a rare population of cells characterized by their multipotency, self-renewal capabilities, and low metabolic and proliferative states. Residing within specialized niches in the bone marrow (BM), HSCs are essential for the continuous production of mature blood cells. To prevent excessive cycling and the accumulation of mutations, HSCs remain quiescent(*1, 2*). Therefore, daily hematopoiesis is predominantly driven by hematopoietic progenitor cells, with minimal contribution from HSCs(*3, 4*). Under stress conditions, however, HSCs respond to increased hematopoietic demands by enhancing the production of downstream progenitors(*5, 6*). Extensive research has explored the biology of HSCs within the BM, particularly examining the effects of cell intrinsic and extrinsic factors on their behavior. Nevertheless, the properties of HSCs outside the BM, especially under long term stress conditions, remain less understood.

Chronic inflammation is a pathological condition characterized by persistent low-grade inflammation with elevated levels of pro-inflammatory cytokines and chemokines. Numerous studies have demonstrated that such inflammation can compromise HSC function in the BM by inducing replicative stress, leading to DNA damage, and stem cell exhaustion(*7–9*). Remarkably, the consequences of chronic inflammation extend beyond diminished stem cell functionality, potentially triggering HSC mobilization and migration to peripheral sites(*10, 11*). This adaptive mechanism, known as extramedullary hematopoiesis (EMH), occurs as hematopoietic cells relocate to organs previously active in fetal hematopoiesis, to sustain blood cell production when BM hematopoiesis becomes insufficient. Spleen and liver are notably most well-documented sites for EMH, featuring unique microenvironments distinct from those of the BM(*12*). EMH is often associated with a myeloid-biased output, a hallmark particularly evident in the context of chronic inflammation and cancer(*13, 14*). Pro-inflammatory cytokines such as interleukin-1β (IL-1β) and interleukin-6 (IL-6), which are usually elevated in pathological and stress conditions, induce proliferation and myeloid differentiation of BM hematopoietic stem and progenitor cells (HSPCs) (*6, 15*). However, the impact of chronic inflammation on HSCs located at the sites of EMH remains poorly understood.

To investigate the effect of chronic inflammation on HSCs located at EMH sites, we utilized a mouse model of chronic multifocal osteomyelitis (CMO), which resembles human autoinflammatory disease chronic recurrent multifocal osteomyelitis (CRMO)(*16*). CMO mice carry a missense mutation, L98P, in the gene *Pstpip2* coding for proline-serine-threonine phosphatase-interacting protein 2 (Pstpip2). The mutation results in misfolding and consequent degradation of the Pstpip2 protein(*17, 18*). Pstpip2 is expressed in immune cells, such as macrophages and neutrophils, but it is undetected in HSCs(*18–20*). Pstpip2 is known to play an anti-inflammatory role by regulating reactive oxygen species (ROS) production and cytokine levels, including IL-1β and IL-6(*18, 19, 21*). Consequently, CMO mice exhibit a progressive sterile auto-inflammatory disorder, with mice being born healthy and developing disease signs at around 7 weeks of age(*22*). Recently, we demonstrated that IL-6 drives an expansion of the HSC pool and diminishes HSC function in the BM, compromising BM hematopoiesis(*20*).

Here, we demonstrate that extramedullary HSCs are not only localized in the blood and spleen of CMO mice but also at sites of inflammation. Using single-cell transcriptomics, we identified *Cd53* as a top differentially expressed gene in extramedullary HSCs. This observation was coupled with the upregulation of MHCII-associated and immunosuppressive genes. Moreover, we showed that extramedullary CD53+ HSCs function as antigen-presenting cells, facilitating the development of regulatory T cells (Tregs) which potentially mitigate chronic inflammation at extramedullary hematopoietic sites. Additionally, we observed that Tregs play a protective role for extramedullary HSCs. Together, our findings unveil a bidirectional interaction between a distinct subset of HSCs and T cells in inflamed tissues during chronic inflammation.

## RESULTS

### CMO mice exhibit increased numbers of functional HSPCs in peripheral blood and spleen

Since hematopoiesis can occur outside of the BM under inflammatory stress(*23–25*), we explored whether our mouse model of sterile chronic inflammation exhibits EMH. We assessed the presence of HSPCs in peripheral blood and spleen of WT and CMO mice and observed increased number of LKS (Lin-c-Kit+ Sca-1+), multipotent progenitors (MPP; LKS CD48+ CD150-) and HSCs (LKS CD48-CD150+) in CMO samples in comparison to WT (Fig. 1, A and B; fig. S1, A to C). These circulating and SP-resident HSPCs were functional, as evidenced by a significant increase in colony numbers observed in cultures established from CMO as opposed to WT cells (Fig. 1, C to E; fig. S1, D to F). Next, we assessed the functionality of extramedullary HSPCs *in vivo* by transplanting blood and splenocytes isolated from WT or CMO mice (Ly5.2) into lethally irradiated Ly5.1 congenic recipients (Fig. 1F; fig. S1G). Consistent with our *in vitro* results, analysis of blood and BM from recipient mice 16 weeks after transplantation revealed increased engraftment in recipients transplanted with CMO blood and splenocytes compared to recipients transplanted with WT control cells (Fig. 1G; fig. S1H). Remarkably, although CMO mice exhibit an expansion of the myeloid lineage(*18, 20*), we did not detect a myeloid bias in blood and spleen of mice transplanted with CMO samples. Instead, we observed enhanced lymphoid production (Fig. 1H; fig. S1I). Altogether, these experiments indicate that CMO mice display increased number of phenotypical and functional HSPCs in circulation and spleen, supporting the existence of EMH in CMO mice.

**Fig. 1.**
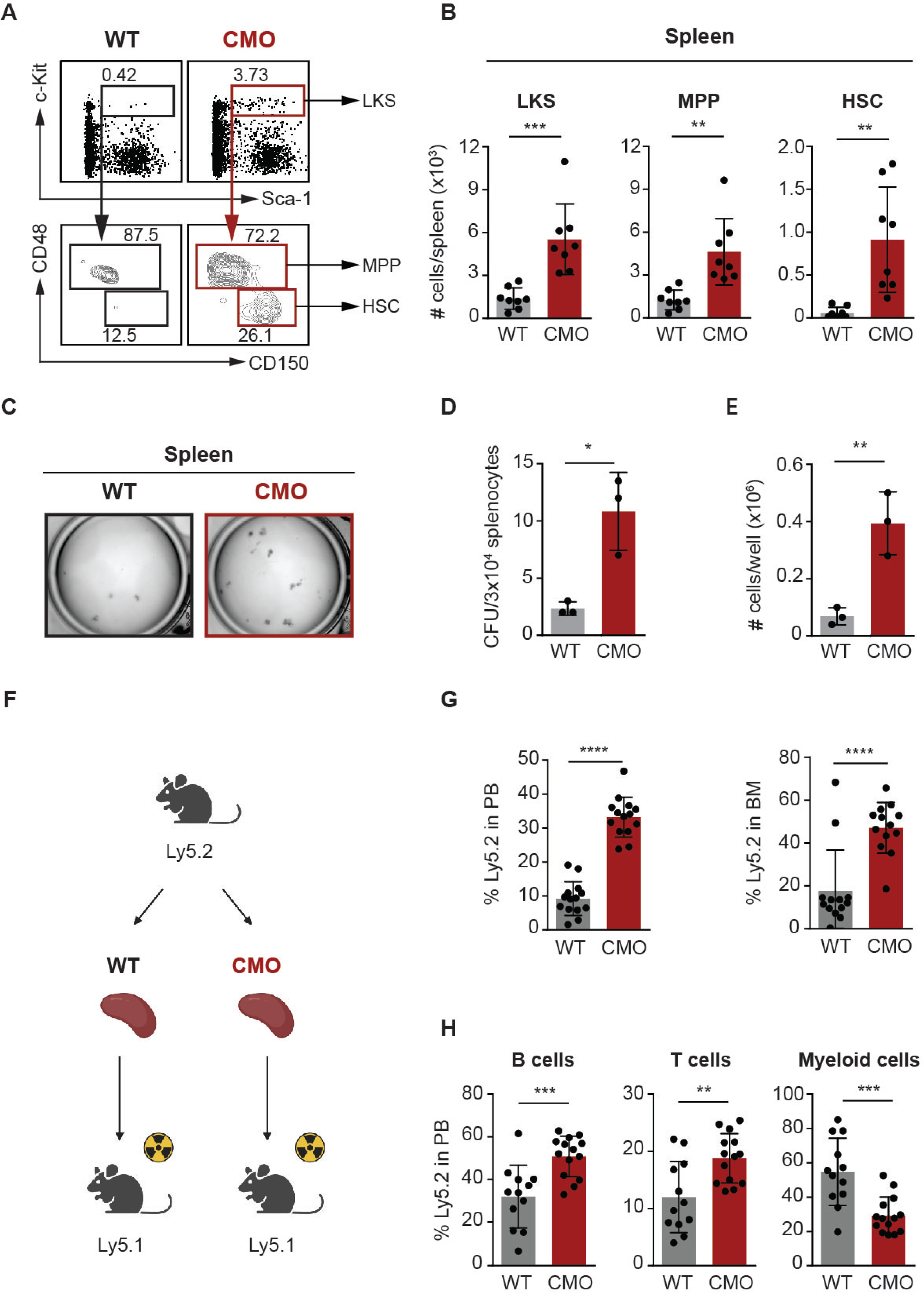
CMO mice show increased levels of functional HSPCs in spleen. (A) Representative flow cytometry plots from 1 WT and 1 CMO spleen. Upper plots indicate c-Kit and Sca-1 levels in lineage-(Lin-) splenocytes. Gates indicate Lin-c-Kit+ Sca-1+ (LKS) cells. Lower plots show CD48 and CD150 expression in LKS, and gates indicate LKS CD48+ CD150-cells (MPPs) and LKS CD48-CD150+ cells (HSCs). Numbers show percentage from parental gates. (B) Quantification of panel A. Y-axes indicate the number of LKS cells, MPPs, and HSCs in WT (gray) and CMO (red) spleens. 8 to 10 mice were included per group in 3 independent experiments. Each mouse is represented by a dot symbol. (C) Representative microscopy images of colony culture assays using MethoCult M3434. Images correspond to day 7 of culture. 3×10^4^ WT (gray) and CMO (red) splenocytes were plated per well. (D-E) Number of colony-forming units (CFU) (D) and cells (E) enumerated in colony culture assays at day 7. Y-axes indicate the numbers per well. At least 3 spleens were used in each condition. (F) Schematic representation of the transplantation setup. 1×10^6^ WT (gray) or CMO (red) of WT or CMO splenocytes were transplanted into the lethally irradiated congenic mice along with 0.5×10^6^ bone marrow support cells. 16 weeks post-transplantation recipients were sacrificed and analyzed. (G) Engraftment of WT (gray) or CMO (red) splenocytes upon transplantation into lethally irradiated congenic recipients. Y-axis indicates the percentage of donor-derived Ly5.2+ cells in peripheral blood (PB, left panel) and BM (right panel) of recipient mice 16 weeks post-transplantation. WT mice are indicated in gray and CMO in red. 12 recipients were used per group in 2 independent experiments, and each dot symbol indicates values per one recipient. (H) Tri-lineage reconstitution analysis in BM of recipient mice 16 weeks after transplantation. Y-axis indicates the percentage of donor-derived Ly5.2+ B, T, and myeloid cells. Each dot indicates values for 1 recipient mouse, and at least 12 mice used in each group. In this figure, data indicate mean ± SD and 2-tailed Student t test was used to assess statistical significance (*P, 0.05, **P, 0.01, ****P, 0.0001).

### CMO mice have ongoing EMH at the sites of inflammation

Since CMO mice exhibit signs of EMH, we next investigated whether this model of sterile chronic inflammation would harbor HSPCs at the sites of inflammation. Thus, we assessed the presence of HSPCs in tails and/or paws, sites commonly inflamed in CMO mice and characterized by inflammatory bone damage, high cytokine levels, and elevated number of granulocytes (*18, 21, 26*). As expected, HSPCs were not present in WT paw, but we detected HSPCs in the inflamed paw of CMO mice (Fig. 2, A and B; fig. S2A). Colony forming assays demonstrated that cell suspensions from WT paws and tails were not able to form colonies, whereas those from CMO paws and tails produced multiple colonies, indicating the presence of functional HSPCs at the inflammatory sites (Fig. 2, C to E; fig. S2, B to D). To further delineate the functionality of CMO paw HSCs *in vivo*, we performed extreme limiting dilution transplantation assays. As depicted in Fig. 2F, CMO HSCs were sorted from paws, mixed with bone marrow support, and transplanted into lethally irradiated congenic recipients. Given the lack of HSCs in WT paws, we compared the engraftment efficacy of CMO paw HSCs with WT BM HSCs. Remarkably, CMO paw HSCs were able to engraft and reconstitute the hematopoietic system of lethally irradiated recipient mice similarly to WT BM HSCs (Fig. 2G). Further, despite the significant increase of granulocytes in paws from CMO mice, CMO paw HSCs exhibited a reduced contribution to the myeloid lineage and a significant increase of lymphocyte production (Fig. 2H). Altogether, our results demonstrate the presence of functional HSCs able to efficiently execute hematopoiesis within the heavily inflamed tissues characteristic of the CMO mice, with a differentiation preference towards the lymphoid lineage.

**Fig. 2.**
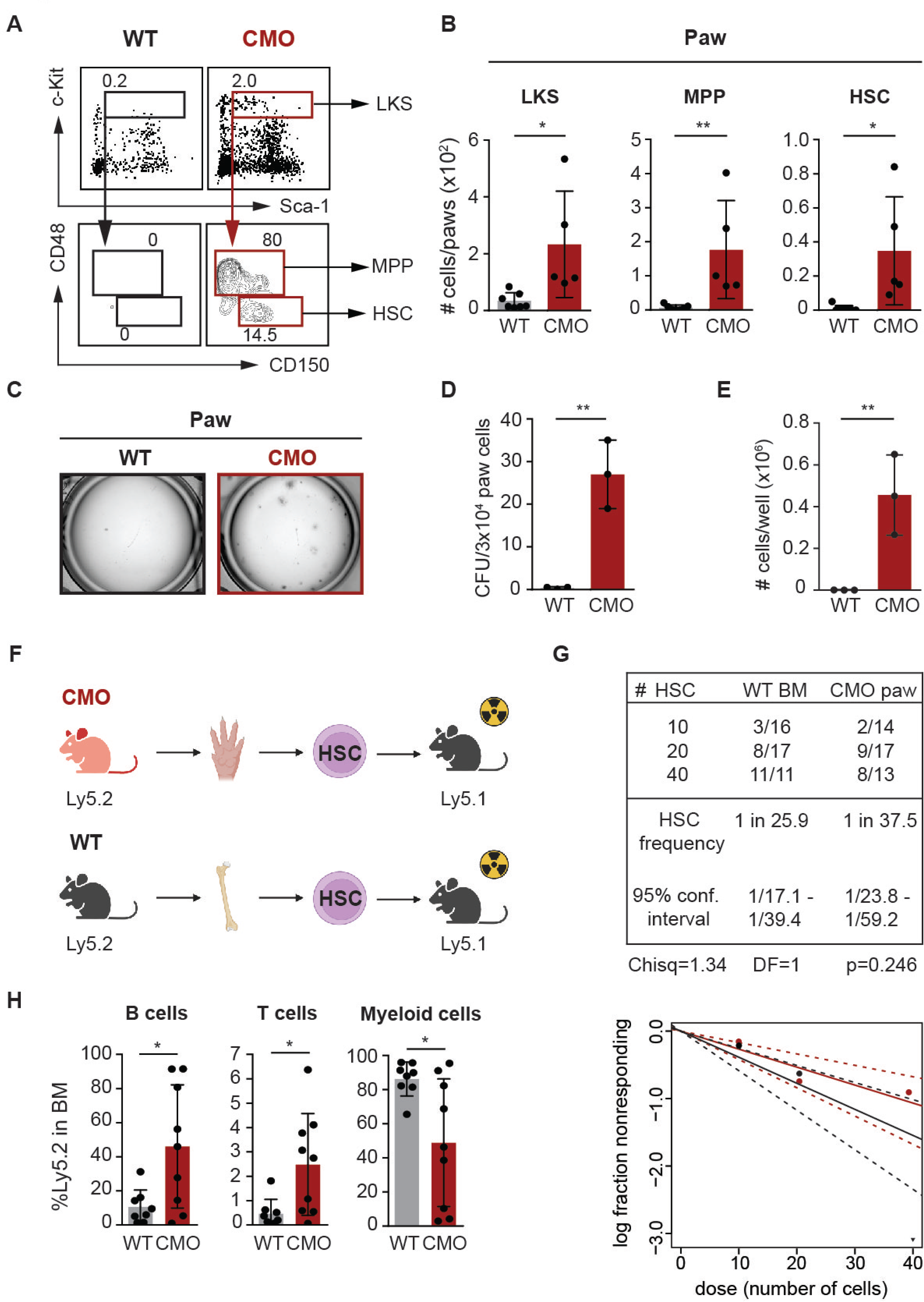
CMO mice exhibit ongoing EMH at the inflamed paws. (A) Flow cytometry plots from 1 WT and 1 CMO paw cell suspension. c-Kit and Sca-1 expression in lineage-(Lin-) cells (upper plots). Rectangles indicate Lin-c-Kit+ Sca-1+ (LKS) cells. Lower plots gated on LKS cells show CD48 and CD150 expression. Boxes illustrate gating for LKS CD48+ CD150-cells (MPPs) and LKS CD48-CD150+ cells (HSCs). Numbers indicate percentage from parental gates. (B) Quantification of distinct cell populations from panel A. Y-axes indicate the number of LKS, MPP, and HSC in cell suspensions from WT (gray) and CMO (red) paws. At least 5 mice were included per group in 2 independent experiments, and each mouse is represented by a dot symbol. (C) Representative microscopy images of colony culture assays after 7 days of culture. 3×10^4^ WT (gray) and CMO (red) paw cells were plated per well using MethoCult M3434. (D-E) Number of colony-forming units (CFU) (D) and cells enumerated in panel (C). Y-axes indicate the numbers per well at day 7. Paw cell suspensions from 3 mice were used in each condition. (F) Schematic representation of the transplantation setup. 10, 20 and 40 WT BM HSCs or CMO paw HSCs were transplanted into lethally irradiated congenic mice along with 0.5×10^6^ BM support cells. 16 weeks post-transplantation recipients were sacrificed and analyzed. (G) Frequency of functional WT BM HSCs and CMO paw HSCs measured by limiting dilution competitive repopulation unit assays and calculated using ELDA online software based on Poisson distribution statistics (Chi-square test; Chisq = 1.34; *P* = 0.246). Graph shows the curve fit of the log fraction of nonresponding mice (solid lines) and confidence intervals (dashed lines) versus the number of mice tested. In logarithmic plot, X-axis indicates the dose of transplanted cells and Y-axis percentages of negative responders. A mouse was counted as responder when engraftment in BM was > 0.1 % and at least 2 out of 3 lineages were reconstituted by donor HSCs. Data compile 3 independent experiments. (H) Tri-lineage reconstitution analysis in BM from responder mice which received 20 WT BM HSCs (gray) or 20 CMO paw HSCs (red). Y-axis indicates the percentage of donor-derived B, T, and myeloid cells. Each dot symbol indicates values for 1 responder mouse. In panels B, D, E, and H data indicate mean ± SD, and 2-tailed Student t test was used to assess statistical significance (*P, 0.05, **P, 0.01).

### Extramedullary HSCs exhibit a unique transcriptional profile

To explore potentially distinct transcriptional profiles of HSCs located in different tissues under chronic inflammatory conditions, we employed single cell RNA sequencing (Sort-seq), a technique which enables the transcriptomic analysis of a low number of cells. As illustrated in Fig. 3A, HSCs were sorted from CMO BM, CMO spleen, and CMO paw, as well as from WT BM, and subjected to Sort-seq. Interestingly, the HSC transcriptional profile was significantly different in the HSC populations analyzed (Fig. 3B), and unsupervised clustering allowed us to identify 4 different HSC identities (cluster 1 to cluster 4) (Fig. 3C). This distribution was defined by specific gene expression patterns in each cluster (Fig. 3D; table S1). Cluster 1 gathered HSCs from WT as well as CMO BM, and enrichment analysis of the genes defining this cluster revealed significant enrichment for transcripts associated with stemness (fig. S3, A to C). Further, we identified two clusters, cluster 2 and 4, which compiled HSCs from WT and CMO BM as well as a smaller proportion of extramedullary HSCs (fig. S3A). These clusters were enriched for the presence of transcripts associated with cell cycle, defining a larger subpopulation of HSCs in the G1/S phase (cluster 2), and a smaller subpopulation of HSCs in the mitotic phase (cluster 4) (fig. S3, D to G). Remarkably, the analysis revealed that HSCs from the CMO spleen and paw predominantly formed a unique cluster, referred here as cluster 3, with minimal overlap to WT and CMO BM HSCs (Fig. 3, C to E). The identification of cluster 3 indicated that HSCs located at the sites of EMH in CMO mice share a unique transcriptional profile, suggesting the existence of unique biological properties and cellular characteristics (Fig. 3E). In fact, analysis of the top differentially expressed genes in this extramedullary cluster revealed enrichment of transcripts associated with cytokine signaling and adhesion/migration (Fig. 3, F and G). Altogether, we observed that HSCs isolated from extramedullary sites in CMO mice share a unique transcriptional profile distinct from BM HSCs, suggesting that inflammation in the extramedullary sites defines the transcriptional properties of these HSCs.

**Fig. 3.**
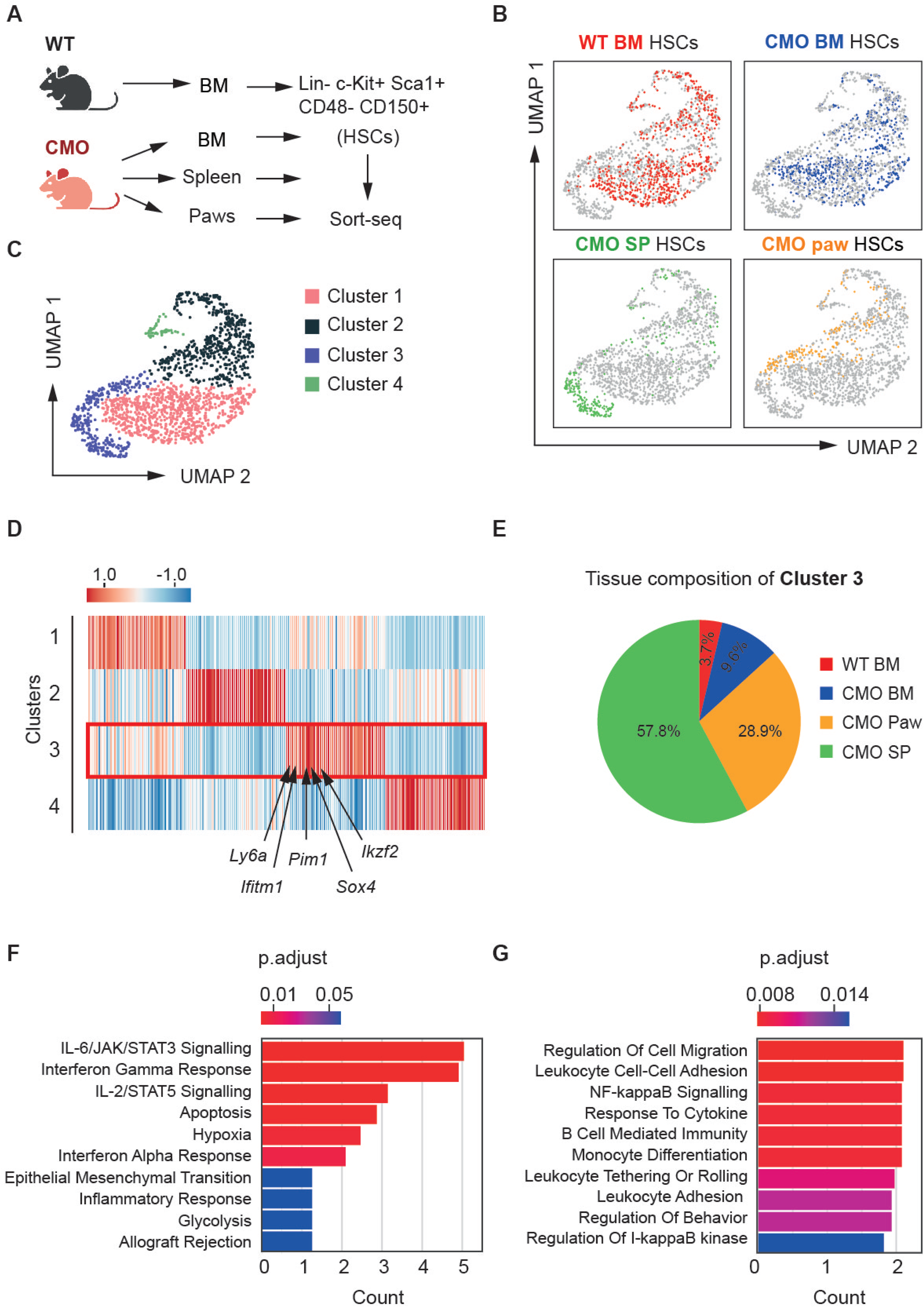
HSCs located in extramedullary sites exhibit a unique transcriptional profile. (A) Graphical representation showing the isolation and sorting of HSCs from different origins. HSCs were defined as Lin-c-Kit+ Sca-1+ CD48- and CD150+ cells and sorted for subsequent sort-sequencing analysis. (B) UMAP plots color-coded for HSCs isolated from the distinct tissues: WT BM HSCs (red), CMO BM HSCs (blue), CMO SP HSCs (green), and CMO paw HSCs (orange). (C) UMAP plot of HSC transcriptomes displaying four clusters identified through unsupervised clustering. (D) Relative gene expression levels of 50 top-ranking marker genes for each identified cluster. Red box highlights cluster 3 and several genes significantly upregulated are indicated. (E) Relative contribution of WT BM, CMO BM, CMO SP and CMO paw HSCs to the cellular composition of cluster 3. (F-G) Relevant results of enrichment analysis of differentially expressed genes by Enrichr (*53*) in cluster 3.

### CD53 expression is increased in extramedullary HSCs and identifies low proliferating and highly efficient stem cells

Next, we further explored the characteristics of HSCs isolated from CMO extramedullary sites. Examination of differentially expressed genes in extramedullary cluster 3 identified *Cd53* as a prominently upregulated gene in HSCs from both spleen and paw of CMO mice (Fig. 4, A and B). We then examined total CD53 protein expression, including intracellular as well as extracellular membrane expression in HSCs from CMO BM, CMO spleen, and CMO paw by flow cytometry. This revealed an upregulation of CD53 levels in CMO spleen and paw HSCs compared to CMO BM HSCs, supporting our Sort-seq results (Fig. 4, C and D). On the other hand, surface expression of CD53 showed different pattern. Not all HSCs in CMO mice had upregulation of CD53 on the cell surface, and in fact, we could divide HSCs into CD53- and CD53+ populations based on the CD53 cell surface levels (fig. S4, A and B). Next, we assessed the levels of Ki-67 proliferation marker in CD53- and CD53+ HSCs from distinct CMO tissues. While no differences were present between CD53- and CD53+ CMO BM HSCs, CD53+ HSCs isolated from extramedullary sites exhibited reduced Ki-67 levels compared to their CD53-HSC counterparts (Fig. 4E). Furthermore, we evaluated the colony-forming capacity of CD53- and CD53+ HSCs isolated from CMO spleen in semi-solid media. Following a 10-day incubation, we observed that CD53+ HSCs exhibited an increased ability to form colonies in comparison to CD53-HSCs (Fig. 4F). To assess the functionality of extramedullary CD53- and CD53+ HSCs isolated from CMO spleen *in vivo*, cells were transplanted in extreme limiting dilution manner (Fig. 4G). 16 weeks upon transplantation, blood analysis demonstrated increased engraftment and hematopoietic reconstitution in recipient mice who received CD53+ spleen HSCs in comparison to recipients transplanted with CD53-spleen HSCs (Fig. 4H). Altogether, we identified a unique set of HSCs located at extramedullary sites, which expresses CD53, exhibits diminished proliferation, and retains increased stem cell properties.

**Fig. 4.**
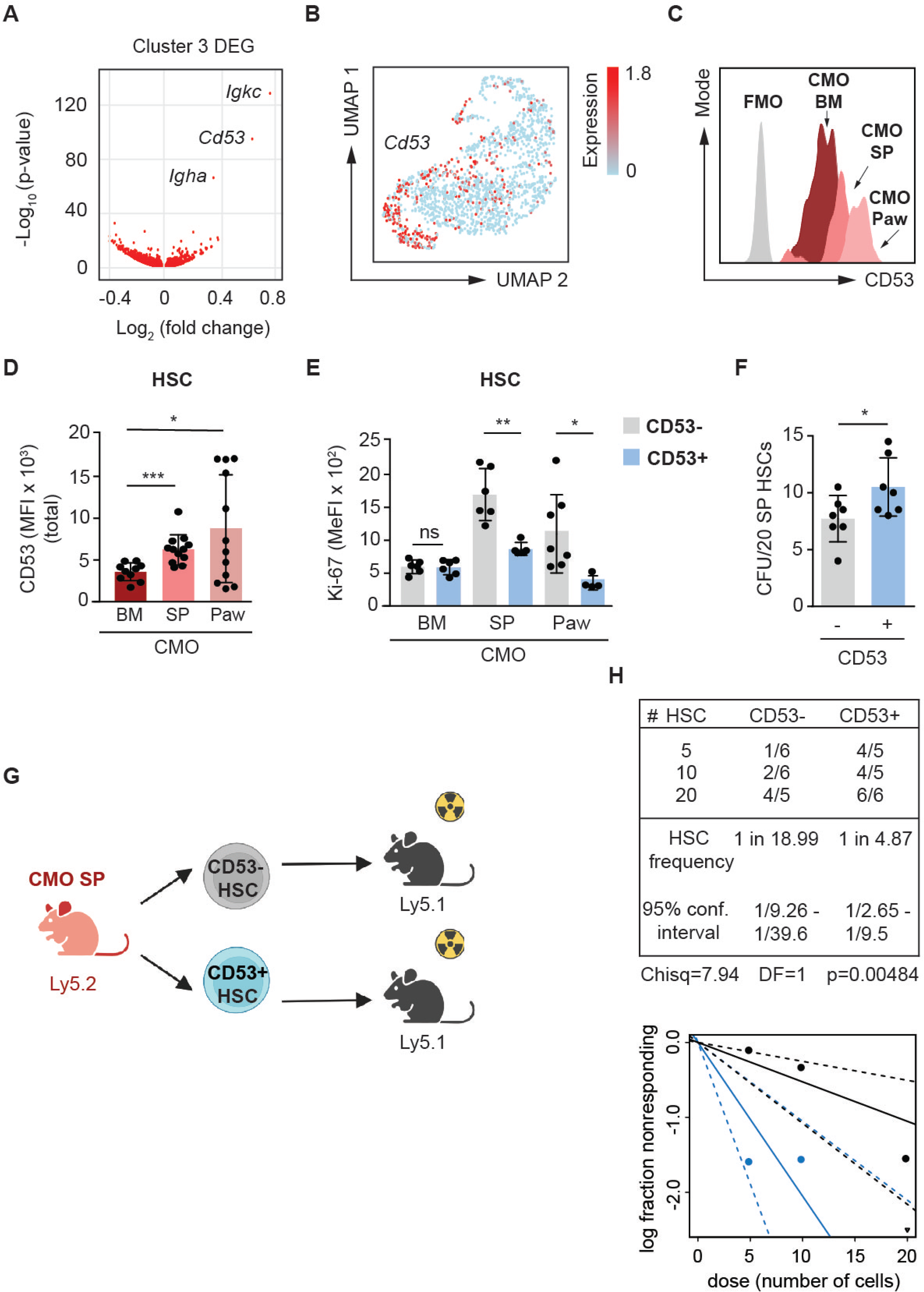
CD53 expression is increased in extramedullary HSCs and identifies low proliferating HSCs with increased engraftment ability. (A) Volcano plot of the most differentially expressed genes in cluster 3 compared to the other identified clusters. (B) UMAP plot displaying relative gene expression of *Cd53* in all HSC clusters. Dark red indicates higher expression levels, while light blue indicates lower expression levels. (C) Representative histogram plot of total (intracellular+surface) CD53 expression in CMO bone marrow (BM), CMO spleen (SP) and CMO paw. Gray peak indicates signal for fluorescence minus one (FMO). (D) Total CD53 levels in HSC (Lin-c-Kit+ Sca-1+ CD48-CD150+) from BM, SP and paw from CMO mice. Y-axis indicates CD53 mean fluorescence intensity (MFI). Each symbol indicates values for one mouse. (E) Ki-67 levels in CD53-HSCs (gray) and CD53+ HSCs (blue) from BM, SP and paw from CMO mice. Y-axis indicates median fluorescence intensity (MeFI) for Ki-67. (F) Quantification of colony forming units (CFU) from CD53-(gray) and CD53+ (blue) HSCs. 20 SP HSCs were plated in Methocult M3434 and the number of colonies was assessed 10 days post-plating. Y-axis indicates number of colonies. Data in panels D, F, and G indicate mean ± SD from at least 2 independent experiments. Each dot indicates values for one biological sample. (G) Schematic representation of the transplantation setup. 5, 10 and 20 CD53- or CD53+ HSCs isolated from CMO SP were transplanted into lethally irradiated congenic mice along with 0.5×10^6^ BM support cells. 16 weeks post-transplantation recipients were sacrificed and analyzed. (H) Frequency of functional CD53- and CD53+ HSCs from CMO SP measured by limiting dilution competitive repopulation unit assays and calculated using ELDA online software based on Poisson distribution statistics (Chi-square test; Chisq = 7.94; P = 0.00484). Graph shows the curve fit of the log fraction of nonresponding mice (solid lines) and confidence intervals (dashed lines) versus the number of mice tested. Logarithmic plot where X-axis indicates the dose of transplanted cells and Y-axis percentages of negative responders. A mouse was counted as responder when engraftment in PB was > 0.01 % and at least 2 out of 3 lineages were reconstituted by donor HSCs. Data show 1 representative experiment. 2-tailed Student t test was used to assess statistical significance in panels D-F (*P, 0.05, **P, 0.01, ***P, 0.001 and ns, not significant).

### Extramedullary HSCs exhibit upregulation of immunosuppressive and MHCII-associated genes

Since our results suggest that extramedullary HSCs may exhibit properties distinct from those of HSCs located in BM, we further explored our transcriptomic data for unique features of extramedullary HSCs. First, we noticed that there was an upregulation in the expression of genes implicated in antigen presentation via MHC Class II molecules in Cluster 3 (Fig. 5A), and accordingly we observed that MHCII surface levels were upregulated in CMO HSCs from spleen and paw compared to BM (Fig. 5B). In addition to the upregulation of MHCII, we noted enhanced expression of immunosuppressive regulatory genes, including genes coding for PD-L1 (CD274) and receptors for IL-10 and TGFβ (Fig. 5C). The collective upregulation of immunosuppressive and MHCII-associated genes in extramedullary HSCs may indicate that these HSCs are trying to counteract the heavy inflammation by acquiring anti-inflammatory properties.

**Fig. 5.**
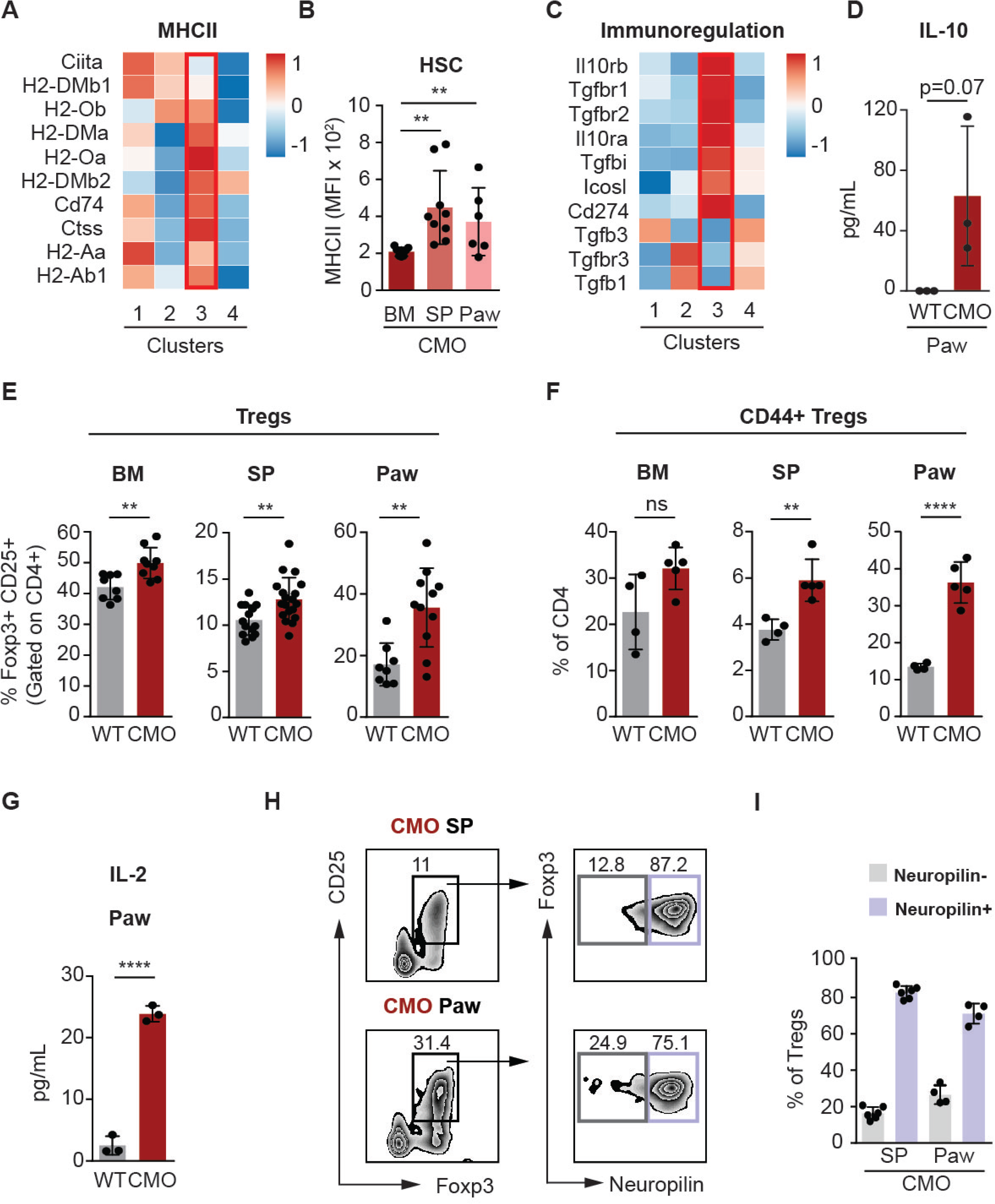
Extramedullary HSCs express high levels of MHCII-associated and immunoregulatory genes, and are surrounded by active regulatory T cells (Tregs) (A) Relative expression of genes associated with MHCII across the distinct HSC clusters. Red box highlights cluster 3 which is composed of extramedullary HSCs. (B) MHCII levels in BM, SP and paw HSCs from CMO mice. Y-axis indicates MHCII mean fluorescence intensity (MFI). Each dot symbol indicates values for 1 mouse. Data indicate mean ± SD from 3 independent experiments. (C) Relative expression of genes associated with immunoregulation across the distinct HSC clusters. Red box highlights cluster 3 which is composed of extramedullary HSCs. (D) IL-10 levels in WT and CMO paw. Y-axis indicates units as pg/mL. Each dot symbol indicates values for 1 mouse. (E-F) Frequency of Tregs (E) defined as CD3+CD4+CD25+Foxp3+ and activated CD44+ Tregs (F) in BM, SP and paw of WT and CMO mice. Y-axes indicate percentage (%) out of CD4+ cells. Each dot symbol indicates values for 1 mouse. Data indicate mean ± SD from 3 independent experiments. (G) IL-2 levels in WT and CMO paw. Y-axis indicates units as pg/mL. (H) Gating strategy for Neuropilin levels in Tregs from SP (upper plots) and paws (lower plots) from CMO mice. Left plots indicate CD25 and Foxp3 levels, and black boxes gate for Tregs. Right plots indicate Foxp3 and Neuropilin expression. Gray boxes indicate Neuropilin negative Tregs and violet boxes Neuropilin positive T regs. Numbers indicate percentages of parental gate. (I) Quantification of panel f. Neuropilin-Tregs (gray) and neuropilin+ Tregs (violet) in CMO SP and CMO paw. Y-axis indicates percentage of Tregs. In this figure, data indicate mean ± SD and 2-tailed Student t test was used to assess statistical significance (**P, 0.01, ****P, 0.0001, and ns, not significant).

### Extramedullary sites in CMO mice exhibit pro- and anti-inflammatory properties

To further characterize the environment in the inflammatory sites, we performed cytokine and chemokine profiling. We observed high levels of chemokines and pro-inflammatory cytokines in extracts from CMO paws (fig. S5, A and B). Nevertheless, this was accompanied by an upregulation of the anti-inflammatory cytokine IL-10 (Fig. 5D). In addition, we noted a significant increase of regulatory T cells (Tregs) across all CMO tissues (Fig. 5E), coupled with increased Treg activation in the spleen and paws, as indicated by the CD44 marker (Fig. 5F). Further, we found elevated IL-2 levels in CMO paws (Fig. 5G), cytokine which is known to enhance T cell proliferation and activation. Despite the increase of Treg in CMO tissues, there was a noticeable overall decline in the frequencies of CD3+ T cells, affecting especially CD8+ T cells (fig. S5C). Additionally, a significant increase in the frequency of CD4+ T cells was observed in the paws of CMO mice, although no notable differences were found in the CD4+ populations within the BM and spleen (fig. S5C). Next, since Tregs can be induced peripherally, we assessed the expression of Neuropilin, which distinguishes thymic-derived Tregs (Neuropilin+) from those induced peripherally (Neuropilin-)(*27*). As anticipated, the majority of Tregs originated in the thymus, while a small fraction appeared to be generated on-site (Fig. 5, H and I). Collectively, our findings indicate that extramedullary HSCs are influenced by a milieu of pro- and anti-inflammatory factors in an environment enriched for the presence of activated Tregs.

### Extramedullary CD53+ HSPCs promote Treg development as a self-protective mechanism

Next, we investigated whether MHCII upregulation preferentially occurs in CD53- or CD53+ HSCs. We observed a significant increase of MHCII expression in CD53+ HSCs compared to CD53-HSCs (Fig. 6, A to C), and this positive correlation was also evident in other splenic populations (fig. S6, A and B). This observation suggests that CD53+ HSCs may act as antigen presenting cells (APCs) and display antigens to T cells at the sites of inflammation, contributing to the development of peripherally-derived Tregs in spleen and paw of CMO mice (Fig. 5, H and I). To address this hypothesis, we sorted Lin-c-kit+ CD11c-CD53+ and Lin-c-kit+ CD11c-CD53-HSPCs from CMO spleen and co-cultured them with naïve T cells isolated from OT-II transgenic mice (Fig. 6D) in the presence of ovalbumin peptide (OVA). After four days, CD53+ HSPCs showed a greater ability to activate naïve T cells compared to CD53-HSPCs, as shown by the increased proliferation of T cells isolated from co-cultures established with CD53+ HSPCs (Fig. 6, E and F) and by the low percentage of anergic T cells (Fig. 6G). Moreover, the viability of T cells was enhanced in the presence of CD53+ HSPCs in comparison to CD53-HSPCs (Fig. 6H). Of note, the enhanced T cell proliferation and viability in CD53+ HSPC co-cultures was similar to that obtained in the presence of professional APCs dendritic cells (Fig. 6 E-H). Interestingly, the co-cultures with CD53+ HSPCs generated higher number of Tregs compared to co-cultures with CD53-HSPCs (Fig. 6I). Next, we investigated the effects of T cells generated in these co-cultures on HSPCs themselves. We observed that CD53+ HSPCs exhibit a higher proportion of c-kit+ cells after 4 days of co-culture with T cells in the presence, but not in the absence, of OVA, suggesting that their interaction with OVA-stimulated T cells in these co-cultures preserves the stem and progenitor pool (Fig. 6J). Collectively, these results suggest that extramedullary CD53+ HSCs may promote the proliferation and viability of T cells and enhance the development of Tregs to preserve the HSPC compartment at the sites of inflammation.

**Fig. 6.**
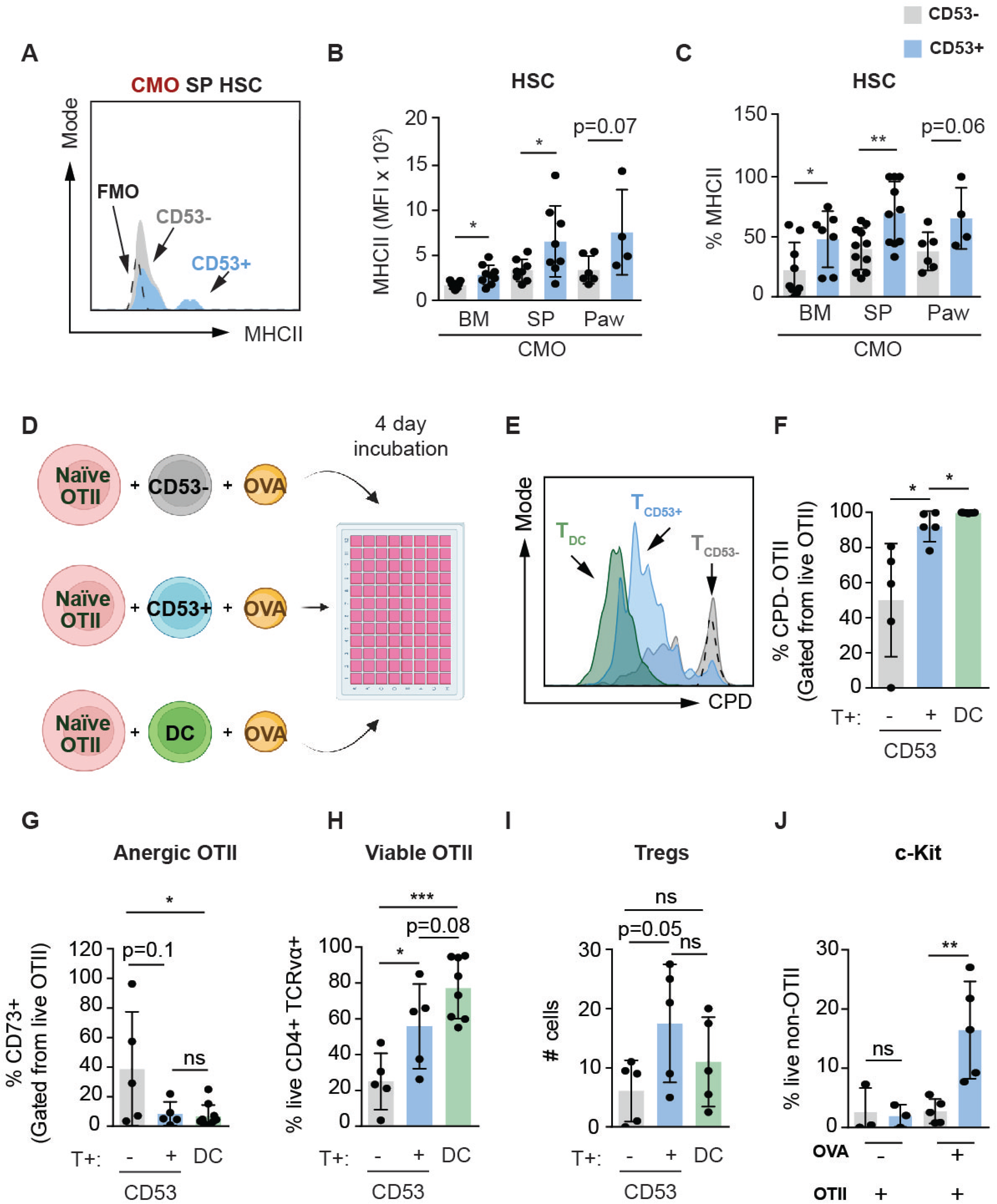
Mutual HSC/Treg effects in extramedullary sites during chronic inflammation. (A) Representative histogram plot of SP HSCs from CMO mice. Y-axis indicates MHCII levels in CD53-(gray) and CD53+ (blue) HSCs. Gray peak indicates signal for fluorescence minus one (FMO). (B-C) Flow cytometric analysis of MHCII levels in CD53-(gray) and CD53+ (blue) HSCs from CMO BM, CMO SP and CMO paw. Y-axis indicate Mean Fluorescence Intensity (MFI) of MHCII signal (B) or percentage (%) of MHCII positive cells (C). (D) Schematic representation of co-culture experiments. (E) Representative histogram plot of T cell proliferation after 4 days in co-culture with dendritic cells (T_DC_; green), CD53+ HSPCs (T_CD53+_; blue), or CD53-HSPCs (T_CD53-_; gray) in the presence of ovalbumin 247-264 peptide (OVA). The negative control (dashed black line) represents naïve CD4+OTII T cells + OVA without antigen presenting cells. X-axis indicates levels of Cell proliferation dye (CPD). (F) Frequency of proliferated CD4+OTII T cells after 4 days of co-culture with CD53-HSPCs (gray), CD53+ HSPCs (blue) and DCs (green) in the presence of OVA. Y-axis indicates percentage of proliferated CD4+OTII cells. (G) Frequency of anergic CD4+OTII T cells after 4 days of co-culture with CD53-HSPCs (gray), CD53+ HSPCs (blue) and DCs (green) in the presence of OVA. Y-axis indicates percentage of anergic CD4+OTII cells. (H) Frequency of viable CD4+OTII T cells after 4 days of co-culture with the indicated cells in the presence of OVA. Y-axis indicates percentage of viable CD4+OTII cells. (I) Number of OTII Tregs after 4 days of co-culture with the distinct cells types and in the presence of OVA. (J) Frequency of c-Kit+ HSPCs after 4 days of co-culture with naïve CD4+ OTII cells. First 2 columns represent negative control co-cultures containing HSPCs and naïve CD4+OTII cells without OVA. The other columns represent co-cultures containing HSPCs, naïve CD4+OTII cells, and OVA. HSPCs were CD53-(gray) or CD53+ (blue).

### Depletion of Tregs in CMO mice results in detrimental effects on extramedullary HSC

To further elucidate the role of Tregs on the HSPC compartment at inflammatory sites, we depleted Tregs in CMO mice using an anti-CD25 antibody (Fig. 7A; fig. S7A). A visualexamination of the anti-CD25- and control-treated mice demonstrated that Treg depletion enhanced the inflammatory phenotype of CMO mice, as observed by the degree of inflammation in paws (Fig. 7B). Since the degree of inflammation in CMO mice correlates with the presence of neutrophils in the inflamed tissues(*28, 29*), we assessed the percentage of neutrophils in BM, spleen and paw of treated mice. Accordingly, a significant increase of neutrophils was observed in the spleen of anti-CD25-treated mice in comparison to control-treated mice (Fig. 7C), however, no significant changes were observed in BM and paw of anti-CD25-treated mice (fig. S7B). Next, we evaluated the frequencies of HSCs following the treatment of CMO mice. We noted a significant reduction in HSC frequency in the spleen, BM, and paw of anti-CD25-treated mice (Fig. 7D; fig. S7C). This decrease could potentially be attributed to the loss of stemness in HSCs, driven by the elevated inflammation and supported by increased Ki-67 expression in CMO spleen HSCs upon anti-CD25 treatment (Fig. 7, E and F). Additionally, CD53 expression significantly decreased post anti-CD25 treatment (Fig. 7G), while Treg depletion did not change CD53 expression in the BM (fig. S7D). Further, we observed a reduction in MHCII levels in CMO spleen HSCs following anti-CD25 treatment (Fig. 7, H and I). Given our hypothesis that Tregs may exert a protective effect on extramedullary HSCs, we transplanted spleen HSCs isolated from anti-CD25- and control-treated CMO mice into lethally irradiated mice (Fig. 7J). The results indicated that untreated spleen HSCs tended to exhibit higher engraftment rates compared to their anti-CD25-treated counterparts (Fig. 7K). Furthermore, we observed a shift from lymphoid to myeloid bias in the anti-CD25-treated HSCs (Fig. 7K). Altogether, these findings suggest that in chronically inflamed sites the interaction of extramedullary HSCs with Tregs may contribute to the maintenance of low proliferating HSCs which express CD53 and MHCII, highlighting a potentially protective role of Tregs in maintaining the extramedullary HSC pool and sustaining HSC activity outside of the bone marrow.

**Fig. 7.**
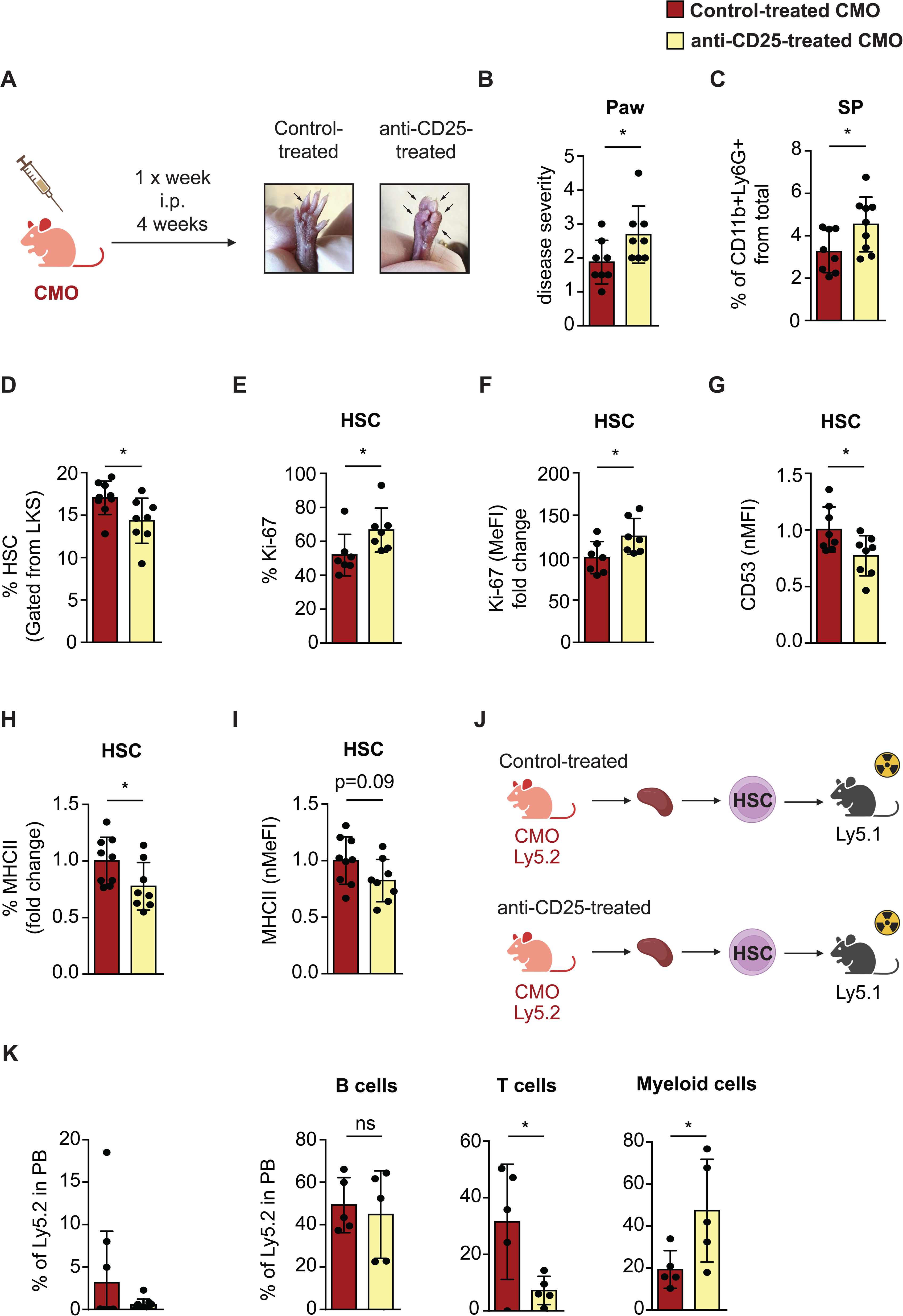
Treg depletion in CMO mice further impairs HSCs. (A) Schematic representation of control and anti-CD25 treatment in CMO mice. Representative pictures illustrate CMO paws upon control treatment (left image) or anti-CD25 treatment (right image). (B) Quantification of panel a. Y-axis indicates the degree of disease severity in paws from control (red) or anti-CD25-treated (yellow) CMO mice. (C) Percentage of CD11b+ Ly6G+ granulocytes in SP of CMO mice. Y-axes indicate percentage of CD11b+ Ly6G+ cells from parental gate. (D) Frequency of HSC in SP isolated from CMO mice. Y-axis indicates percentage (%) from parental LKS gate. (E) Frequency of Ki-67+ in SP HSC from CMO mice. Y-axis indicates percentage (%) from parental HSC gate. (F) Ki-67 levels in SP HSCs from CMO mice. Y-axis indicates median fluorescence intensity (MeFI) for Ki-67. (G) Flow cytometric analysis of total CD53 expression in SP HSC from CMO mice. Total CD53 expression is indicated as a fold change from non-treated (red) CMO group. Y-axis indicates CD53 normalized mean fluorescence intensity (nMFI). (H) Frequency of MHCII+ HSC in SP of CMO mice. Y-axis indicates percentage (%) from parental HSC gate. (I) MHCII levels in SP HSCs from CMO mice. Y-axis indicates normalized median fluorescence intensity (nMeFI) for MHCII. (J) Schematic representation of the transplantation setup. 10 CMO SP HSCs from control- or anti-CD25-treated mice were transplanted into lethally irradiated congenic mice along with 0.5×10^6^ BM support cells. 16 weeks post-transplantation recipients were bled, and engraftment was analyzed. (K) Engraftment of CMO SP HSCs. Y-axis indicates the percentage of donor-derived Ly5.2+ cells in peripheral blood (PB, left panel). 9-10 recipients were used per group. (L) Tri-lineage reconstitution analysis in PB of recipient mice 16 weeks after transplantation. Y-axes indicate the percentage of donor-derived Ly5.2+ B, T, and myeloid cells. In this figure, data indicate mean ± SD from at least 2 independent experiments. All animals included in Fig. 7 were 16-week-old CMO mice. Each dot indicates values for one biological sample. 2-tailed Student t test was used to assess statistical significance (*P, 0.05, and ns, not significant).

## DISCUSSION

During the last decade, we have learned many aspects relevant to BM HSC maintenance and fate, such as their location in the BM niche, transcriptomic profile, metabolic needs, and lineage bias. Nevertheless, the properties and hallmarks of HSCs located outside of the BM remain less understood. In case of inefficient BM hematopoiesis, adult HSCs can exit the BM niche and migrate to alternative hematopoietic sites, establishing EMH. For instance, extramedullary HSCs can be detected in spleen, kidney, and liver during benign hematological conditions, neoplastic disorders, as well as during inflammatory stress. Here, we employ the CMO mouse model, characterized by the presence of sterile chronic inflammation, which demonstrated an abundance of circulating HSPCs in blood and the existence of EMH. We also report that, in addition to the classical tissues associated with EMH, non-classical extramedullary sites, such as the inflamed tails and paws of CMO mice, can also host functional HSPCs. Previously, it was reported that myeloid progenitors can be accommodated in inflamed joints in mice suffering from systemic inflammation(*30*). In the present manuscript, using *in vitro* cultures as well as *in vivo* transplantation assays we demonstrated that not only hematopoietic progenitors but also stem cells are present in inflamed tissues, where HSCs and immune cells co-habit in an inflammatory environment, and potentially influence each other. Since patients that exhibit EMH show ongoing hematopoiesis in classical as well as non-classical extramedullary sites(*31*), we made use of our unique CMO murine model to investigate the properties of HSCs in extramedullary sites and explore their relationship with the local inflammatory environment.

In the present study, we profiled HSCs from WT BM and from CMO BM, spleen, and paw to identify unique HSC subpopulations and properties related to extramedullary HSCs. We determined that extramedullary HSCs share a similar gene expression profile marked by high expression of CD53, a member of the tetraspanin superfamily(*32, 33*). However, while CD53 is uniquely expressed within the immune compartment, its role in HSC biology remains largely unknown. Recent studies have shown that CD53 expression is associated with a quiescent state in both human and murine HSCs(*34, 35*). Accordingly, we observed that extramedullary CD53+ HSCs are less proliferative than CD53-HSCs. Nevertheless, while Greenberg and colleagues reported that CD53 upregulation induces quiescence in stressed HSCs through the DREAM complex, our scRNA-sequencing data did not show upregulation of DREAM complex-related genes (data not shown), pointing towards another mechanism responsible for the decreased proliferation observed in CD53+ HSCs. Alternatively, the involvement of the DREAM complex could be caused by the use of distinct models, ranging from endogenous CD53 expression to CD53 knockout mice(*35*), or by activation of the DREAM complex at the protein level despite no changes in RNA. Previous studies have demonstrated that CD53 supports migration and mobilization of immune cells(*36, 37*). In line with these reports we observed that extramedullary HSCs have upregulated genes associated with cell migration and mobilization, such as *Selp*, *Sell*, *Pecam1*. Nevertheless, despite transcriptomic profiling suggesting that these extramedullary HSCs possess a migratory phenotype, it remains unknown whether they migrated from the BM to the sites of inflammation, or if they are being produced at the sites of inflammation and are ready to migrate to other tissues. Future research should explore the migratory properties of extramedullary HSCs.

Our results provide two major discoveries that change our understanding of EMH and provide evidence for a mutual relationship between HSPCs and T cells at chronically inflamed extramedullary sites. First, we demonstrate that a subset of HSPCs located at inflammatory sites act as APCs, promoting T cell proliferation, survival, and the development of Tregs, overall coordinating a mechanism trying to regulate inflammation at extramedullary sites. Notably, these extramedullary HSPCs were characterized by the elevated co-expression of CD53 and MHCII, molecules found in close proximity in the plasma membrane of human B cells and dentritic cells(*38*). Previous studies have shown that antigen-presenting BM HSPCs can activate naïve CD4+ T cells(*39*). However, our findings indicate that extramedullary CD53+ HSPCs act as APCs and facilitate the production of Tregs. This is consistent with reports identifying a sub-population of CD53+ megakaryocytes which is able to activate T cells(*40*), altogether suggesting a general role for CD53 in APC regulation. Importantly, not all Tregs in the inflamed sites were thymic-derived Tregs, and there was a fraction of peripherally-induced Tregs. We suggest that CD53+ HSCs contribute to the generation of these peripherally-induced Tregs, and speculate that this peripheral induction is an additional attempt to further modulate inflammation at the inflamed sites. In addition, we show that Tregs protect extramedullary HSCs during chronic inflammation, preserving stem cell properties and supporting hematopoiesis at the extramedullary sites. While the expression of MHCII in all BM HSCs was previously reported(*39*), our results reveal that only a subset of extramedullary HSCs express MHCII. Mechanistically, we observe that MHCII expression in HSPCs triggers distinct immune responses in BM and extramedullary sites. Haas and colleagues reported that BM HSPCs interact with CD4+ T cells to promote HSPC proliferation, differentiation, and exhaustion of aberrant HSPCs during transformation, aiming to eliminate transformed HSPCs and prevent leukemia onset(*39*). In contrast, since BM hematopoiesis is inefficient under chronic inflammation(*20*), we propose that Tregs protect extramedullary HSCs, which support blood cell production at extramedullary sites. This HSC-protective mechanism is in line with previous reports highlighting the vulnerability of HSCs to excessive inflammatory signals(*7, 15*).

While the expression of MHCII on BM HSCs in steady state was reported previously as a mechanism to eliminate transformed HSPCs from the hematopoietic system and prevent transformation, here we provide additional insights into the role of MHCII expression of HSCs during inflammation. We suggest that the interaction between HSCs and Tregs in the inflammatory sites acts as a novel regulatory mechanism aiming to protect the extramedullary HSCs from the hostile inflammatory environment. This highlights the importance of preserving a small extramedullary HSC pool during diseases which leads to ineffective BM hematopoiesis, such as in the case of CMO.

## MATERIALS AND METHODS

### Animals

12- to 25-week-old CMO (Pstpip2 ^cmo/cmo^) and aged-matched C57BL/6J mice were employed. At this age CMO mice displayed visible symptoms of the disease such as swollen paws, tail kinks, and developed EMH. OT-II x Rag1 KO x Foxp3^DTR^ mice and OT-II x Rag1 KO mice (used as FMO controls) were culled at 6 to 12 weeks of age(*41–43*). For the transplantation assays, congenic strains Ly5.1 and Ly5.2 C57BL/6J were used. Mice were maintained under specific pathogen free conditions in the animal facility of the Institute of Molecular Genetics of the CAS. All experiments were approved by the ethical committee of the institute (approval numbers: AVCR 7141-2022 SOV II).

### Flow cytometric analysis and HSC sorting

WT and CMO mice were sacrificed by cervical dislocation, long bones (femurs and tibias), spleen, and paws were isolated. Long bones and paws were processed to single cell suspension by crunching using pestle and mortar. Spleens were processed using syringe pistons. Single cell suspensions were filtered and red blood cells were lysed using ACK for 5 min at room temperature (RT). Next, cells were labelled with fluorescent-dye conjugated antibodies listed in Table S2 and analyzed on Symphony instrument (BD Biosciences, san Jose, CA, USA).

For HSC sorting, tissues were processed as described above using a two-step method. Initially, the lineage positive fraction was tagged using biotinylated lineage-specific antibodies. These cells were then subjected to anti-biotin magnetic beads separation (Miltenyi Biotec, Bergisch Gladbach, Germany), following the guidelines provided by the manufacturer using a MACS separator. Subsequently, the lineage negative (Lin-) fraction was stained with various antibodies to label HSCs. An Influx instrument (BD Biosciences, San Jose, CA, USA) was utilized for sorting the HSCs using a defined strategy (Lin-, c-Kit+, Sca-1+, CD48-CD150+)(*44*). During sorting and flow cytometric analyses, either Hoechst 33258 or Zombie dye UV was included in the cell suspensions to identify and exclude non-viable cells. Antibodies were purchased from BD Biosciences (San Jose, CA, USA), eBioscience (San Diego, CA, USA) or BioLegend (San Diego, CA, USA) (see Table S2). Data were obtained using Diva software (BD Biosciences, San Jose, CA, USA) and analyzed using FlowJo software (Tree Star Incorporation, Ashland, OR, USA).

### Blood transplantation assays

Peripheral blood was collected from 15-week-old WT and CMO mice expressing Ly5.2. Red blood cells were lysed using the ACK buffer. Following the lysis, an equivalent of 800 μL was injected via tail vein into each lethally irradiated congenic mouse (Ly5.1) together with 0.5×10^6^ Ly5.1 support BM cells.

### Splenocyte transplantation assays

Spleens from Ly5.2 WT and CMO C57BL/6J mice were processed with syringe pistons to obtain single cell suspensions. 1×10^6^ splenocytes were counted and then mixed with 0.5×10^6^ Ly5.1 BM support cells before being intravenously transplanted into lethally irradiated Ly5.1 recipients. PB and BM analyses were performed 16 weeks post-transplantation. To distinguish between donor-derived and support cells, cells were labeled with antibodies against Ly5.1 and Ly5.2. Lineage-specific antibodies (B220, CD3, CD11b, and Gr1) were utilized to assess the reconstitution of B-cells (B220+), T-cells (CD3+), and myeloid cells (CD11b Gr1+).

### HSC extreme limiting dilution transplantation assays

For HSC limiting dilution transplantation assays, WT and CMO C57BL/6J mice expressing Ly5.2 were used, while congenic Ly5.1 C57BL/6J mice served as recipients. Donor mice used in these assays were typically 12-25 weeks old. Various doses of HSCs (5, 10, 20, and 40), characterized as LKS CD48− CD150+, were sorted and then transplanted intravenously along with 0.5×10^6^ WT BM (Ly5.1+) support cells. Prior to transplantation, recipient mice underwent lethal irradiation. Analyses of PB and BM were conducted 16 weeks after transplantation. To differentiate between donor-derived and support cells, samples were labeled with antibodies against Ly5.1 and Ly5.2. Lineage-specific antibodies (B220, CD3, CD11b, and Gr1) were used to evaluate the reconstitution of B-cells, T-cells, and myeloid cells. A recipient mouse was considered successfully engrafted if a determined % of the cells were Ly5.2+ (figure legends indicate the specific cut off in each experiment) and contributed to at least two out of three lineages assessed. The frequency of engrafted HSCs was determined using ELDA online software(*45*), which employs Poisson statistics and the method of maximum likelihood to estimate the proportion of negative recipients in a limiting dilution assay.

### Single cell sort, sample preparation, and single cell transcriptomic SORT-seq

One WT and six symptomatic CMO C57BL/6J mice were utilized for single-cell SORT sequencing(*46*). BM HSCs from WT and CMO mice were isolated and sorted directly into 384-well microplates containing well-specific barcoded primers using an Influx instrument (BD Biosciences, San Jose, CA, USA). SP and paws from five CMO mice were processed, and their HSCs were also sorted directly into 384-well microplates. SP and paw cells from the five mice were pooled together, with each tissue type assigned to its own plate. In total, WT BM HSCs were distributed across two plates, CMO BM HSCs across another two plates, and CMO SP and paw HSCs were each allocated to a single plate. The plates were sealed and stored at −80°C before being shipped to Single Cell Discoveries (The Netherlands) for sequencing. Synthetic RNA control sequences (ERCC-00002 to ERCC-00171) were included in the sequenced libraries. The DNA libraries were paired-end sequenced on an Illumina Nextseq™ 500, high output, with a 1×75 bp Illumina kit (read 1: 26 cycles, index read: 6 cycles, read 2: 60 cycles). During sequencing, read 1 was assigned 26 base pairs and was used to identify the Illumina library barcode, UMI (6bp) and cell barcode (8bp). Read 2 was assigned 60 base pairs and used to map to the reference genome with STARSolo 2.7.10b. Mapping and generation of count tables were automated using the STARSolo 2.7.10b aligner.

### Single cell SORT sequencing analysis

Obtained reads were mapped to GRCm38 genome (Ensembl annotation version 98) (*47*). Quality control was performed to filter out low-quality cells. Cells with fewer than 1,000 unique molecular identifiers (UMIs) were excluded from further analysis. Cells with more than 10% of reads mapping to mitochondrial genes were removed to mitigate potential biases from stressed or dying cells. Secondary data analysis was conducted using BIOMEX software(*48*). Following quality control, data were normalized using log-normalization and scaled. Principal component analysis (PCA) was performed on all features, followed by dimensionality reduction using Uniform Manifold Approximation and Projection (UMAP) in two-dimensional space. Graph-based clustering from the Seurat package(*49*) was applied to identify cell populations, resulting in five distinct clusters visualized on the UMAP plot. Ranked top markers from obtained clusters were acquired in a two-step manner. First, differential analysis was performed comparing each cluster against all other clusters where unique upregulated markers were assigned, followed by ranking of genes using a product-based meta-analysis(*50*). In addition, we performed a pair-wise differential analysis of cluster 4 versus all other clusters using limma package(*51*) and ranked the results by log_2_ fold change. Heatmap analysis was performed using heatmaply package (version 0.15.2) (*52*) with cluster-averaged gene expression data that has been auto-scaled for visualization.

### Colony culture assays

Murine colony culture assays were performed using Methocult GF M3434 (Stemcell Technologies, Vancouver, BC, Canada). 3×10^4^ splenocytes and 100 μL blood were plated after cell blood lysis using ACK. 3×10^4^ paw cells and 3×10^4^ tail cells were sorted based on viability and plated. For the CD53 HSC colony cultures, splenocytes from CMO mice were isolated and stained for Lin-, c-Kit+, Sca-1+, CD48-, CD150+ along with CD53. HSCs were sorted as CD53- and CD53+, and subsequently, 20 CD53-HSCs or 20 CD53+ HSCs were plated. In all cultures, colonies were counted and cells harvested after 7-10 days of *in vitro* culture. Pictures of the colonies were obtained on Apotome microscope.

### Intracellular CD53 staining

To assess total CD53 expression in HSCs, cells from BM, spleen, and paw were isolated and processed into a single-cell suspension as described above. Subsequently, the cells were stained with HSC-specific surface markers (Lin-, c-Kit+, Sca-1+, CD48-CD150+) for 30 minutes at 4°C. After staining, the cells were washed twice with PBS containing 2% FBS and fixed using the BD Cytofix/Cytoperm kit (BD Biosciences) for one hour at 4°C. Following fixation, cells were washed with washing buffer and stained for CD53 according to the manufacturer’s instructions. Samples were analyzed by flow cytometry using a Symphony instrument (BD Biosciences, san Jose, CA, USA).

### Ki-67 staining in HSC

BM, spleen, and paw were isolated and processed into a single-cell suspension as described above. HSCs were stained using the following surface markers (Lin-, c-Kit+, Sca-1+, CD48-CD150+) for 30 minutes at 4°C. Next, the cells were washed twice with PBS containing 2% FBS and fixed using the BD Cytofix/Cytoperm kit (BD Biosciences) for one hour at 4°C. Following fixation, cells were washed and stained for Ki-67 according to the manufacturer’s instructions. Samples were analyzed by flow cytometry using a Symphony instrument (BD Biosciences, san Jose, CA, USA).

### Flow cytometric analysis of Tregs

BM, spleen, and paw cells were isolated and processed as described above. First, cells were stained for 30 min at 4°C with surface markers CD3, CD4, CD8, CD25 (to identify T cells) and with Neuropilin (to differentiate between thymic-derived and peripherally-induced T cells). Subsequently, cells were washed twice with PBS containing 2% FBS and fixed/permeabilized using the eBioscience™ Foxp3/Transcription Factor Staining Buffer Set, according to manufacturer’s instructions. After fixation and permeabilization, the Foxp3 antibody was added, and the cells were incubated overnight at 4°C. Next day, samples were washed and analyzed on a Symphony instrument (BD Biosciences, san Jose, CA, USA).

### Co-cultures

HSPCs (Lin-c-kit+ CD11c-CD53- and Lin-c-kit+ CD11c-CD53+) and dendritic cells (DCs; CD11c+MHCII+) were sorted into 96-well U-bottom plates. Each well contained 200 µL of RPMI supplemented with 10% heat-inactivated Fetal Calf Serum (FCS, Gibco), penicillin/streptomycin (100 U/mL, Sigma), sodium pyruvate (1.5 mM, Gibco), and MEM non-essential amino acids (1x, ThermoFisher). Prior to the cell sorting, OTII T cells (OT-II x Rag1 KO x Foxp3^DTR^) were isolated from skin draining and mesenteric lymph nodes mechanistically using 40µm cell strainer. Isolated cells were depleted of unwanted fraction using CD4+ T cell Isolation Kit (Miltenyi) to enrich OTII T cells. Next, enriched OTII T cells were labelled with Cell Proliferation Dye eFluor 670 (eBioscience) according to the manufacturers protocol, stained for cell sorting and sorted to the wells containing HSPCs or DCs in a ratio of 2000 HSPCs/DCs to 4000 naïve CD4+ OTII T cells. OVA peptide (OVA 323-339, InvivoGen) was introduced to stimulate the co-cultures. As a negative control, (OVA 257-264) was added to wells containing HSPCs/DCs and naive CD4+ OTII T cells. Cells were incubated at 37°C and 5% CO2 for 4 days. T cell proliferation, Treg numbers, and HSPC properties were assessed on the fourth day of co-culture.

### Cytokine levels measurement

WT and CMO paws were crunched with a mortar and pestle in PBS, and supernatants were collected and used without freezing. 31 cytokines, chemokines, and growth factors were measured using the Bio-Plex Pro Mouse Chemokine Panel 31-Plex #12009159 kit, according to the protocols of the manufacturer. Data acquisition was made using Bio-Plex 200 System (Luminex, Luminex Corporation, Technology Blvd, Austin Texas, USA) and software Bio-Plex Manager 6.0 (Bio-Rad Laboratories, Inc., 2000 Alfred Nobel Drive, Hercules, CA, USA). Samples were measured under two distinct sensitivity modes (low RP1 value for low sensitivity and high RP1value for high sensitivity).

### CMO mice anti-CD25 treatment

12-week-old CMO mice were treated with InVivoMAb anti-mouse CD25 (IL-2Rα) (cat. #BE0012, BioXcell) over a period of 4 weeks. Mice received a weekly intraperitoneal injection of 0.4 mg per mouse. After 4 weeks, mice were sacrificed, and their BM, spleen, and paws were harvested and processed into single-cell suspensions for flow cytometry analysis.

### Disease severity assessment

Disease severity in CMO mice was assessed by visually inspecting each hind paw, with each inflamed finger receiving a score of 0.5 points. Each cumulative score represents the total disease severity for an individual mouse.

### Statistical analysis

Statistical significance for indicated data sets was determined using two-sided, unpaired Student’s t-test. P-values <0.05 were considered statistically significant. Poisson statistics and the method of maximum likelihood to estimate the proportion of negative recipients was employed in extreme limiting dilution assays.

## Supplementary Materials

**This PDF file includes:**

Figs. S1 to S7

Table S1 and S2

Supplementary references

## Author contributions

MA-J conceived and designed the study; MK, SG, JB, KV, NP, and SR carried out experiments; MK and MA-J performed data analysis and interpretation; MK, JB, DF, TB and MA-J contributed to experimental design; MK, MM, JR and KR performed scRNA-seq data analysis; MK and MA-J wrote the manuscript. All authors revised the final manuscript.

## Data and materials availability

All data needed to evaluate the conclusions in the paper are present in the paper and/or the Supplementary Materials. Sort-seq data are accessible at the following link: https://www.ebi.ac.uk/biostudies/arrayexpress/studies/E-MTAB-14287?key=012f7d76-a976-407c-a090-5c503d75fde3.

---

**Supplementary Figure 1.**
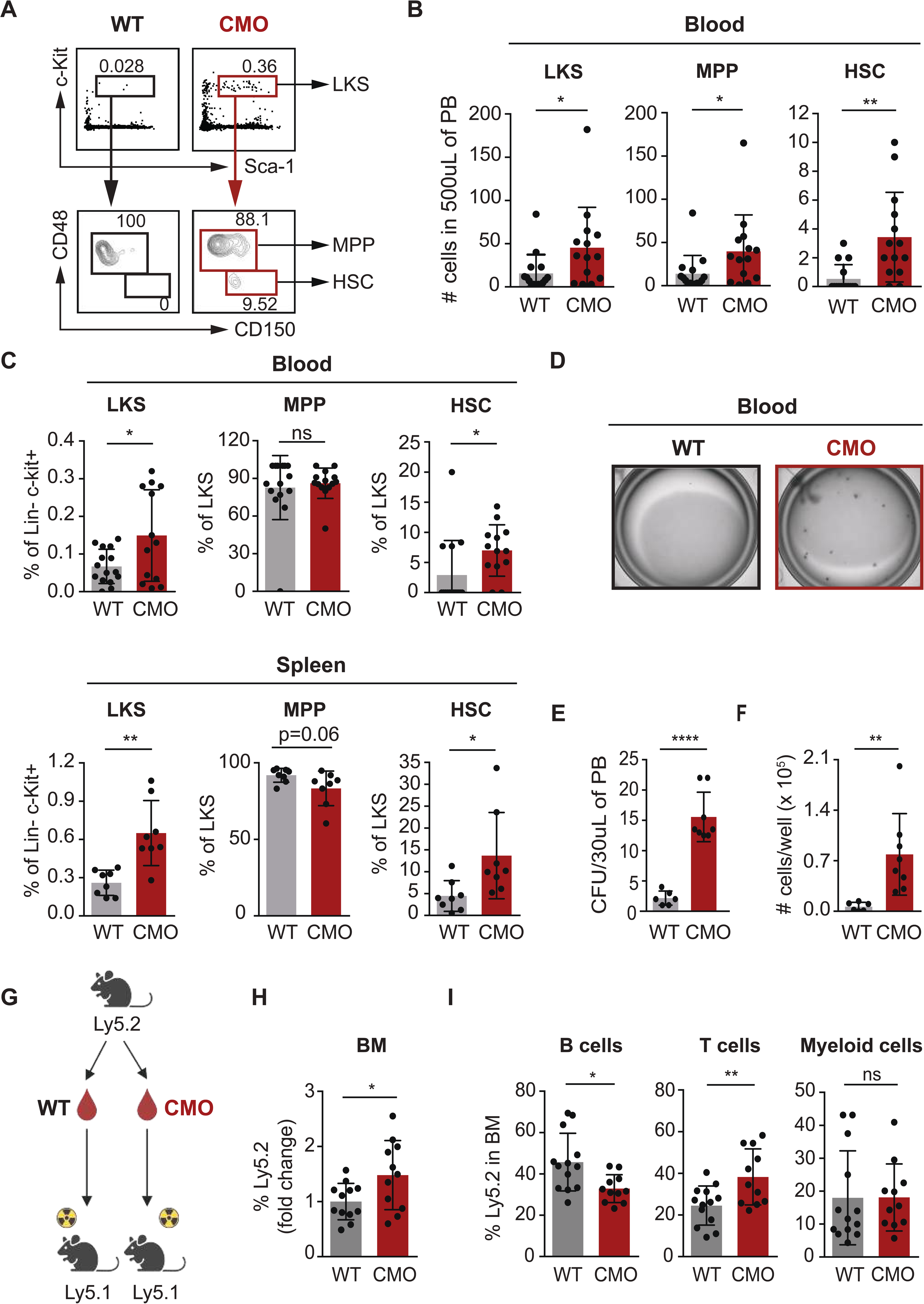
CMO mice exhibit increased numbers of functional HSPCs in peripheral blood. **(A)** Representative flow cytometry plots from blood isolated from 1 WT and 1 CMO mouse. Upper plots illustrate c-Kit and Sca-1 expression in lineage-(Lin-) cells and the rectangle gates for Lin-c-Kit+ Sca-1+ (LKS) cells. Lower plots indicate CD48 and CD150 expression in LKS, and boxes illustrate gating for LKS CD48+ CD150-cells (MPPs) and LKS CD48-CD150+ cells (HSCs). Numbers indicate percentage from parental gates. **(B)** Quantification of panel a. Y-axes indicate the number of LKS, MPP, and HSC in 500 µL of peripheral blood (PB) from WT (gray) and CMO (red) mice. At least 8 mice were included per group, and each mouse is represented by a dot symbol. **(C)** Frequency of LKS, MPP, and HSC in blood and SP from WT (gray) and CMO (red) mice. Y-axes indicate percentage (%) from parental gate. Each dot indicates values for 1 mouse. All animals included were 16 to 20 weeks old. Data indicate mean ± SD from at least 2 independent experiments, and 2-tailes Student t test was used to assess statistical significance (*P, 0.05, **P, 0.01). **(D)** Representative microscopy images of colony culture assays using MethoCult M3434. A total of 30 µL of PB from WT (gray) and CMO (red) mice were plated per well. Images correspond to day 7 of culture. **(E-F)** Number of colony-forming units (CFU) (E) and cells (F) enumerated in panel c. Y-axes indicate the numbers per well at day 7. PB from at least 6 mice in 3 independent experiments was used in each condition. Each mouse is represented by a dot symbol. **(G)** Schematic representation of blood trans-plantation assays. Cells present in 1000 µL of PB from WT or CMO mice (Ly5.2) were transplanted into lethally irradiated Ly5.1 recipient mouse along with 0.5×106 BM support cells (Ly5.1). **(H)** Quantification of engraftment 16 weeks post-transplantation. Y-axes indicate percentage of WT (gray) and CMO (red) donor-derived Ly5.2+ cells in PB and BM. Engraftment is indicated as fold change from WT group. At least 11 animals were included in each group. Each dot indicates values for 1 animal. **(I)** Lineage reconstitution analysis 16 weeks after transplantation in BM from recipients transplanted with blood from WT (gray) and CMO (red) mice. Y-axis indicates the percentage of donor-derived Ly5.2+ B cells, T cells, and myeloid cells. Each dot indicates values for 1 recipient mouse. At least 11 recipients were used in each group. All animals included in Figure 1 were 12 to 25 weeks old. Data indicate mean ± SD from at least 3 indepen-dent experiments, and 2-tailed Student t test was used to assess statistical significance (*P, 0.05, **P, 0.01, ****P, 0.0001, and ns, not significant)

**Supplementary Figure 2.**
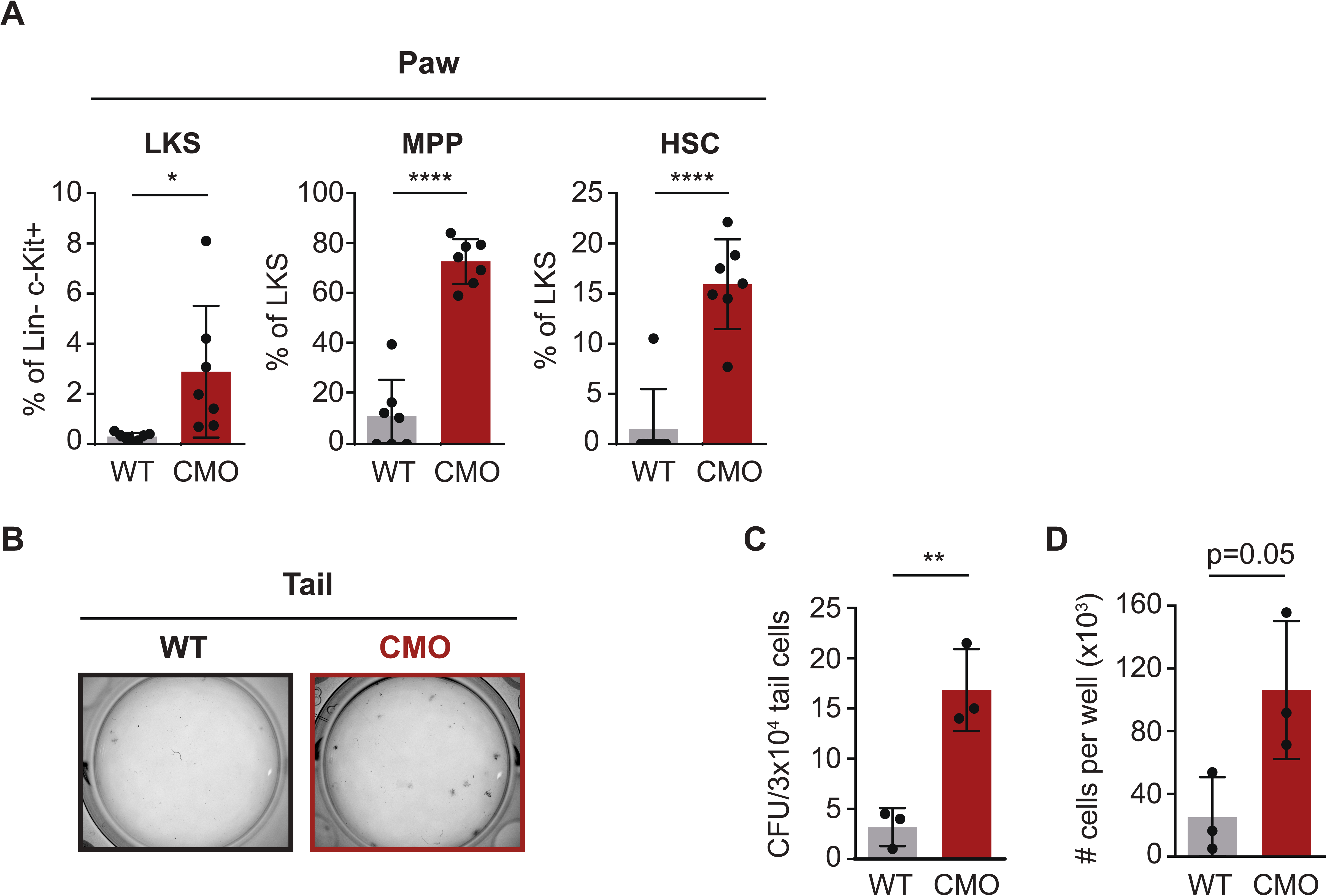
Presence of functional HSPCs in inflamed paws and tails. **(A)** Frequency of LKS (Lin-c-Kit+ Sca1+), MPP (Lin-c-Kit+ Sca1+ CD48+CD150-) and HSC (Lin-c-Kit + Sca1+ CD48-CD 150+) from paw. Y-axis indicates percentage (%) from parental gate in paw from WT (gray) and CMO (red) mice. Each dot indicates values for 1 mouse. All animals included were 16 to 20 weeks old. Data indicate mean ± SD from at least 3 independent experiments, and 2-tailed Student t test was used to assess statistical significance (*P, 0.05, ****P, 0.0001). **(B)** Representative microscopy images of colony culture assays after 7 days of culture. 3×103 WT (gray) and CMO (red) cells from tail were plated per well using MethoCult M3434. **(C)** Enumeration of panel b. Number of colony forming units (CFU) from cells isolated from WT (gray) and CMO (red) tails. Each dot symbol indicates values for one mouse. 2-tailed Student t test was used to assess statistical significance (**P, 0.01). **(D)** Number of cells in cultures from panel b. Y-axes indicate numbers of cells per well at day 7. Tail cell suspensions from 3 mice were used in each condition.

**Supplementary Figure 3.**
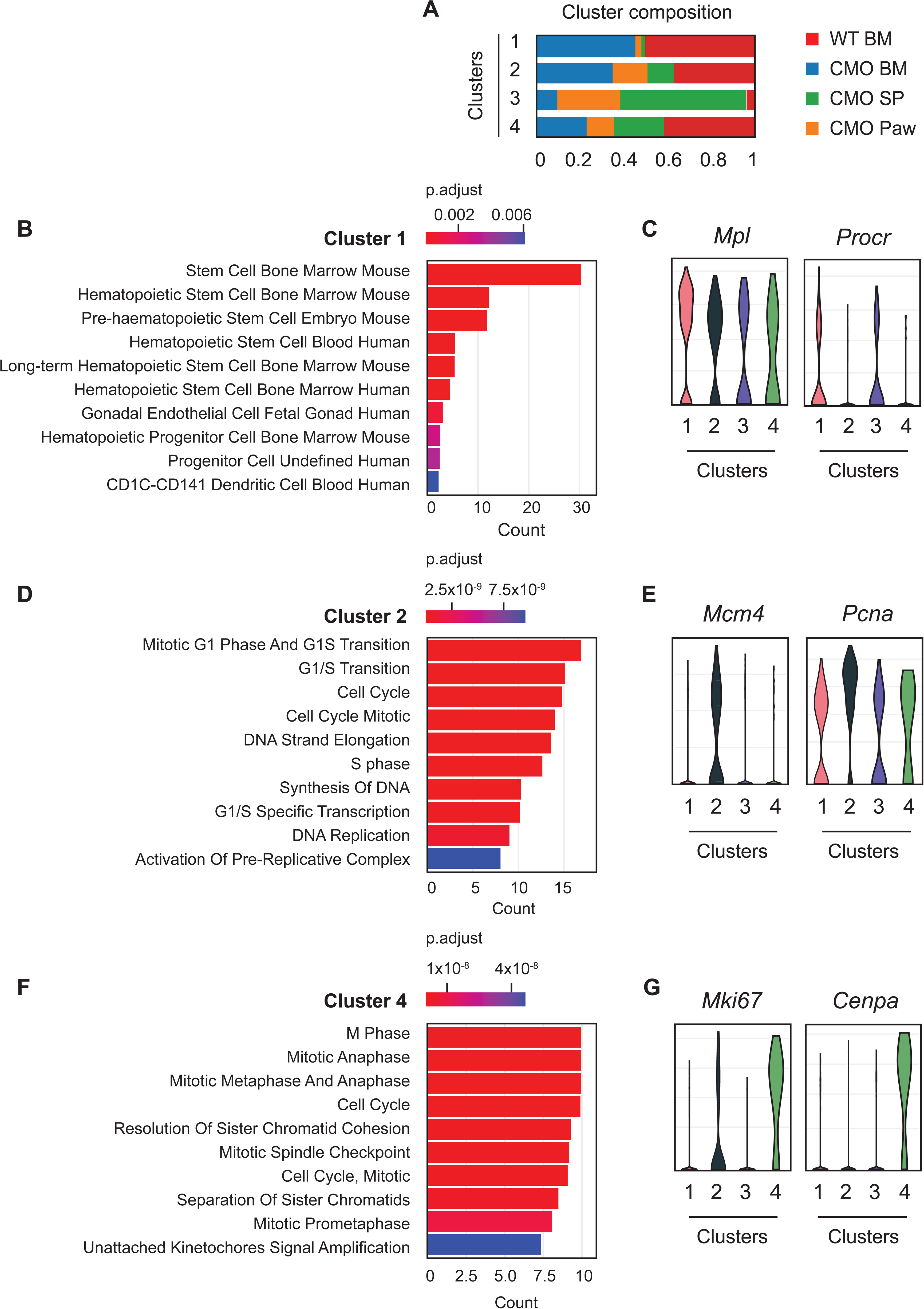
Identification and classification of distinct HSCs clusters. **(A)** Relative contribution of WT BM, CMO BM, CMO SP and CMO paw HSCs to the cellular composition of each cluster. Y-axis indicates the cluster number. X-axis indicates the relative contribution of the individual HSC samples to each cluster. **(B)** Relevant results of enrichment analysis of differentially expressed genes by Enrichr 1 in cluster 1. **(C)** Violin plots representing the expression of Mpl and Procr genes in cluster 1. **(D)** Relevant results of enrichment analysis of differentially expressed genes by Enrichr 1 in cluster 2. **(E)** Violin plots representing the expression of Mcm4 and Pcna genes in cluster 2. **(F)** Relevant results of enrichment analysis of differentially expressed genes by Enrichr 1 in cluster 4. **(G)** Violin plots representing the expression of Mki67 and Cenpa genes in cluster 4.

**Supplementary Figure 4.**
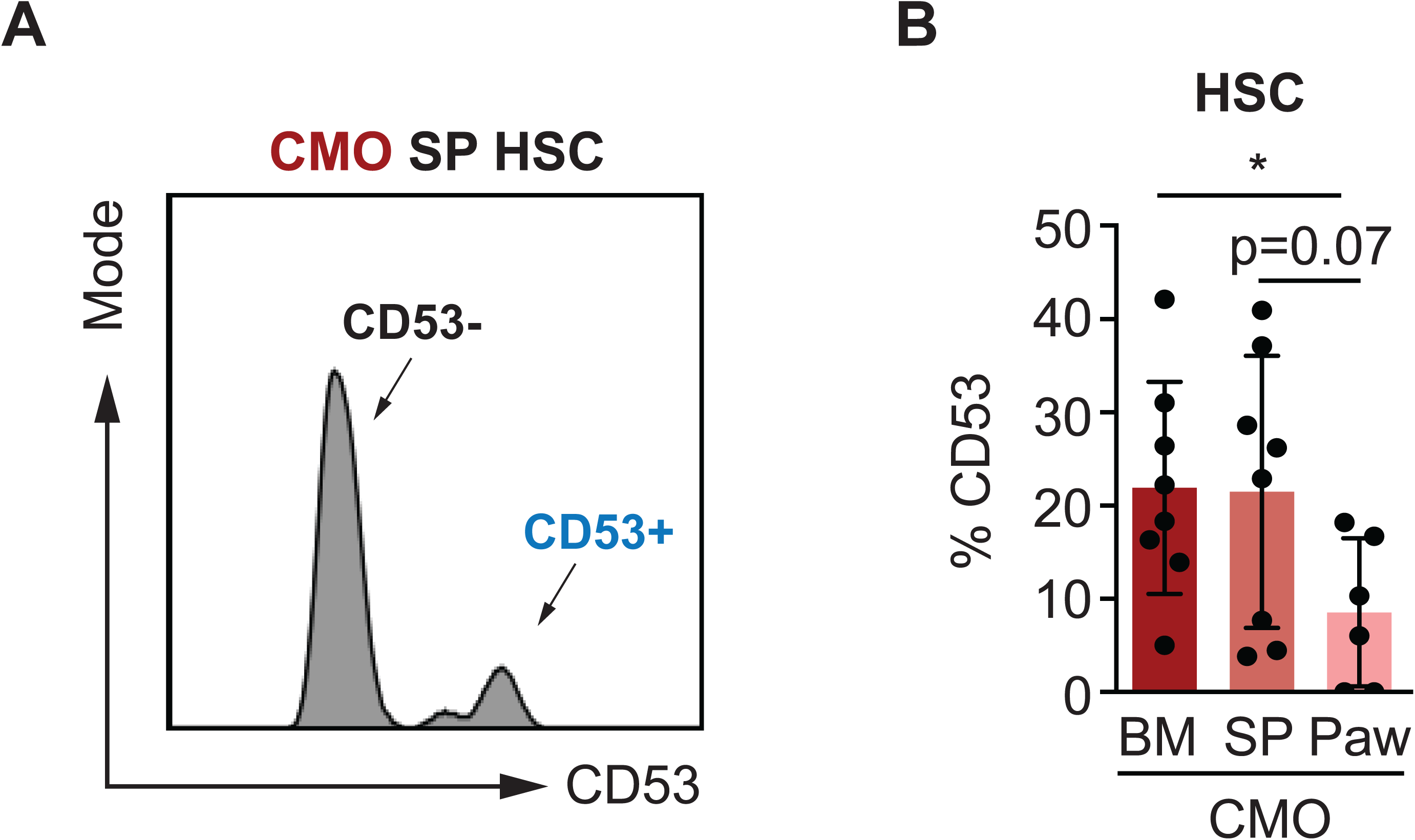
CD53 and Ki-67 labeling in HSCs. **(A)** Representative histogram plot of surface CD53 in CMO SP HSCs. Arrows indicate a subset of CD53- and a CD53+ HSCs. **(B)** Frequency of CD53 in BM, SP and paw HSCs from CMO mice. Y-axis indicates percentage (%) of CD53 in the cell surface of HSCs. All animals included were 12 to 15 weeks old. Data indicate mean ± SD from 3 independent experiments. 2-tailed Student t test was used to assess statistical significance (*P, 0.05, **P, 0.01, and ns, not signifi-cant).

**Supplementary Figure 5.**
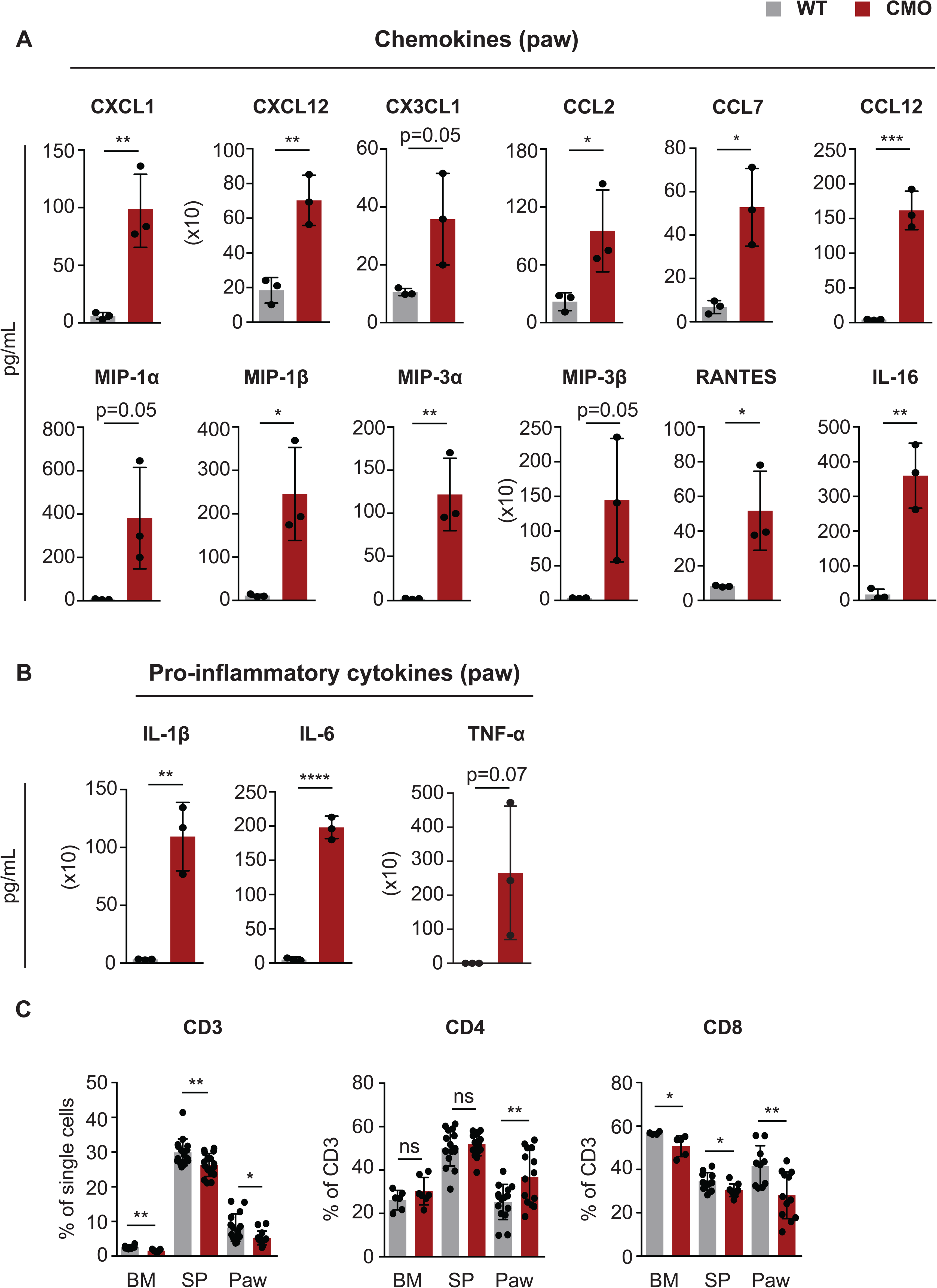
Pro- and anti-inflammatory environment in CMO paws. **(A-B)** Chemokine and cytokine profiling by BioPlex in CMO paw. Each dot symbol indicates values for 1 mouse. Data indicate mean ± SD from 1 independent experiments. **(C)** Frequency of CD3, CD4 and CD8 in BM, SP and paw HSCs from CMO mice. Y-axis indicates percentage (%) of CD3, CD4 or CD8 from parental gate. Each dot symbol indicates values for 1 mouse. Data indicate mean ± SD from 2 and more independent experiments. All animals included were 16 to 25 weeks old. 2-tailed Student t test was used to assess statistical significance (*P, 0.05, **P, 0.01, ****P, 0.0001).

**Supplementary Figure 6.**
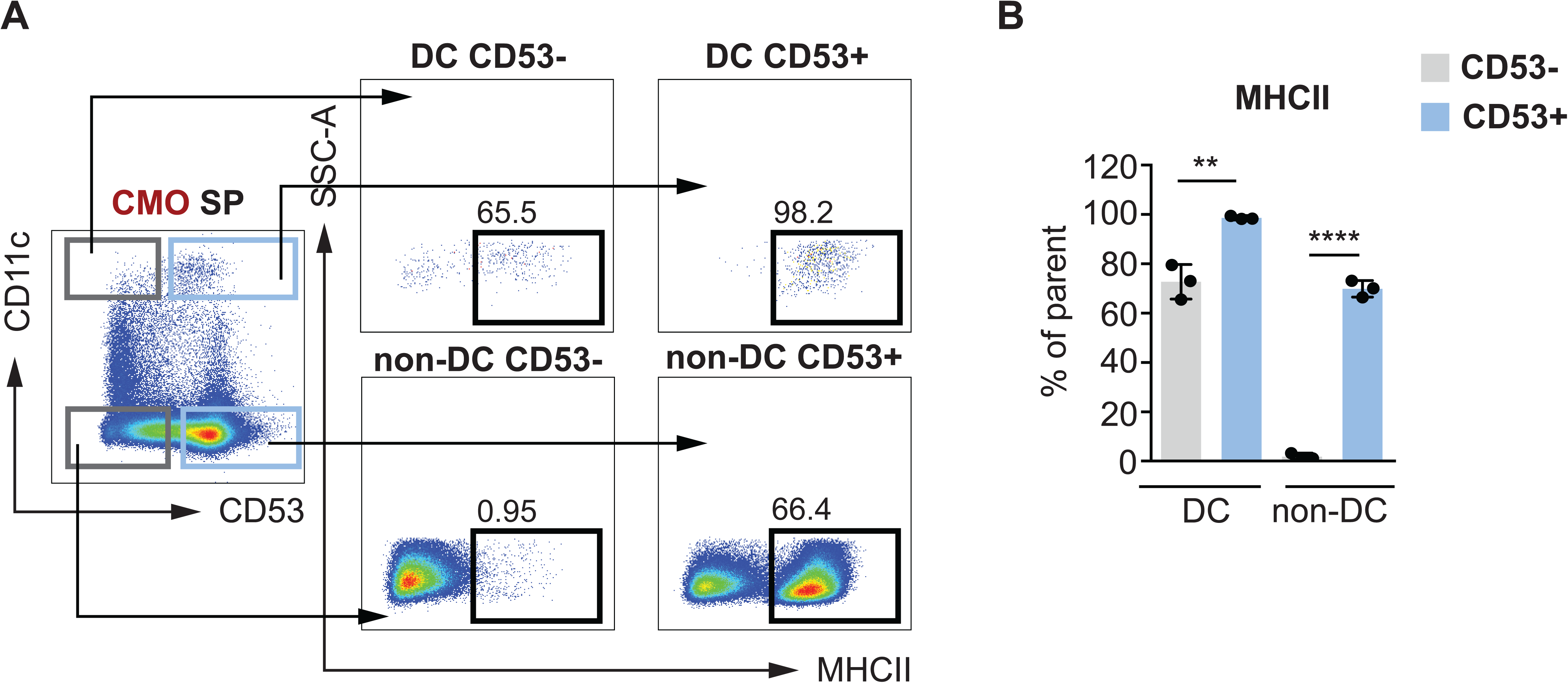
CD53+ cells contain higher levels of MHCII. **(A)** Representative gating strategy for MHCll in CMO SP. Splenocytes were gated according to CD11c and CD53 levels. Upper left box indi cates CD53-dentritic cells (DC CD53-), upper right box CD53+ CD11c+ cells (DC CD53+), lower left box CD53-CD11c-(non-DC CD53-), and lower right box CD53+ CD11c-(non-DC CD53+). The rest of the dot plots indicate MHCll levels in the distinct populations. Black boxes indicate MHCll+ cells. Numbers indicate percentages from parental gates. (B) Frequency of MHCl lexpres-sion across in DC and non-DC pop ulations from CMO SP. CD53-cells are indicated in gray and CD53+ in blue. Each dot symbol indicates values for 1 mouse. Data indicate mean ± SD from 1 experiment. 2-tailed Student t test was used to assess statistical significance (**P, 0.01, ****P, 0.0001).

**Supplementary Figure 7.**
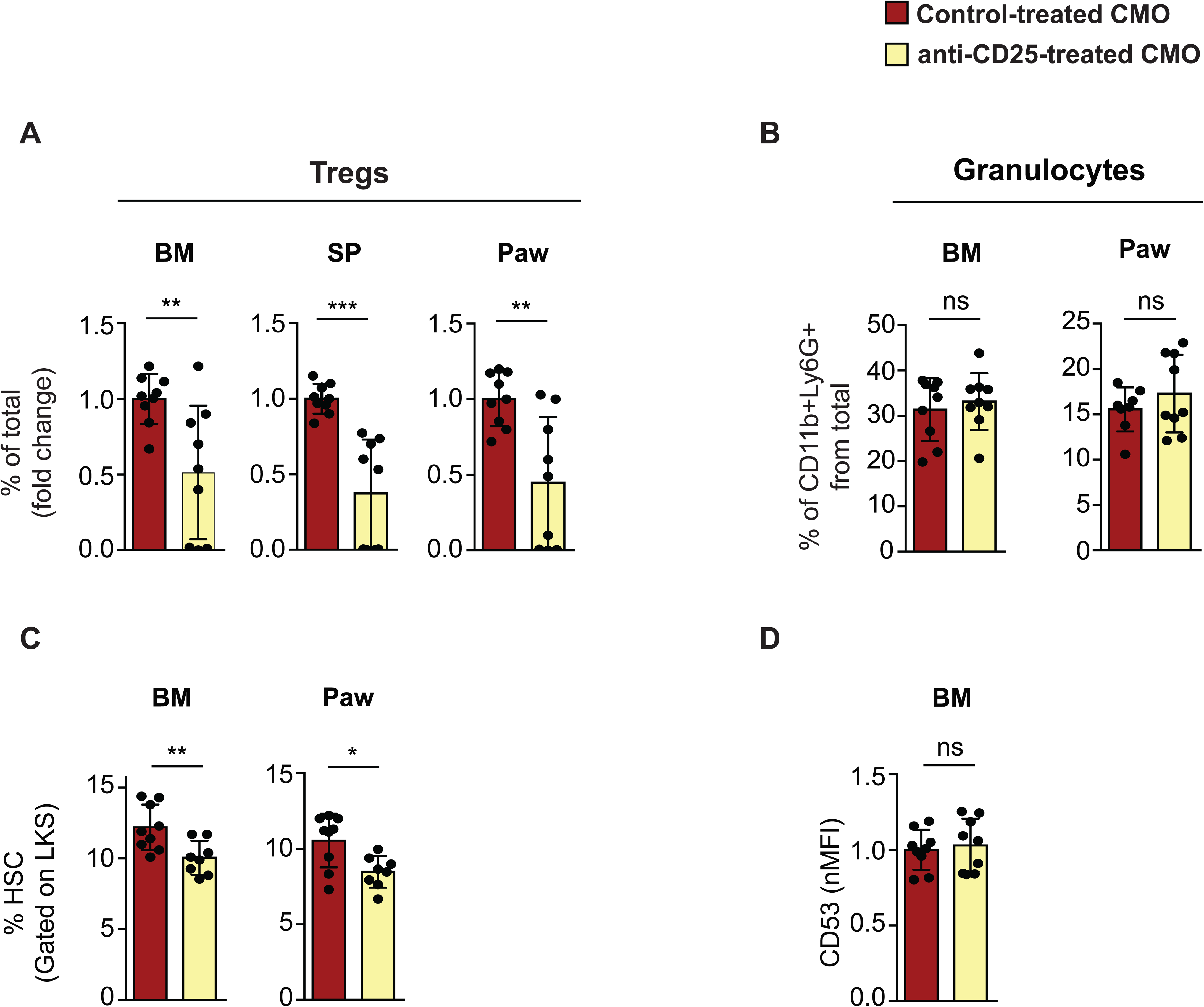
Treg depletion effect on BM and paw HSC. **(A)** Frequency of CD25+Foxp3+ Tregs in non-treated (red) and anti-CD25 treated (yellow) CMO mice in BM, SP and paw. Y-axis indicates percentage (%) of Tregs from total cells and is calculated as a fold change from non-treated (red) CMO group. X-axis indicates whether mice received (yellow) or not (red) anti-CD25 antibody treatment. **(B)** Frequency of CD11b+ Ly6G+ granulocytes in non-treated (red) and anti-CD25 treated (yellow) CMO mice in BM and paw. Y-axis indicates percentage (%) of granulocytes from parental gate. X-axis indicates whether mice received (yellow) or not (red) anti-CD25 antibody treatment. **(C)** Frequency of HSC (Lin-c-Kit+ Sca1+ CD48-CD 150+) from control (red) or anti-CD25 (yellow) CMO treated mice in BM and paw. Y-axis indicates percentage (%) from parental LKS gate. X-axis indicates whether mice received (yellow) or not (red) anti-CD25 antibody treatment. **(D)** Flow cytometric analysis of total CD53 expression in HSC from CMO BM. HSCs were isolated from non-treated (red) and anti-CD25-treated (yellow) CMO mice. Total CD53 expression is indicated as a fold change from non-treated (red) CMO group. Y-axis indicates CD53 mean fluorescence intensity (MFI). X-axis indicates whether mice received (yellow) or not (red) anti-CD25 antibody treatment. Each dot indicates values for 1 mouse. Data indicate mean ± SD from 3 independent experiments. 2-tailed Student t test was used to assess statistical significance (ns, not significant).

**Table S1.**
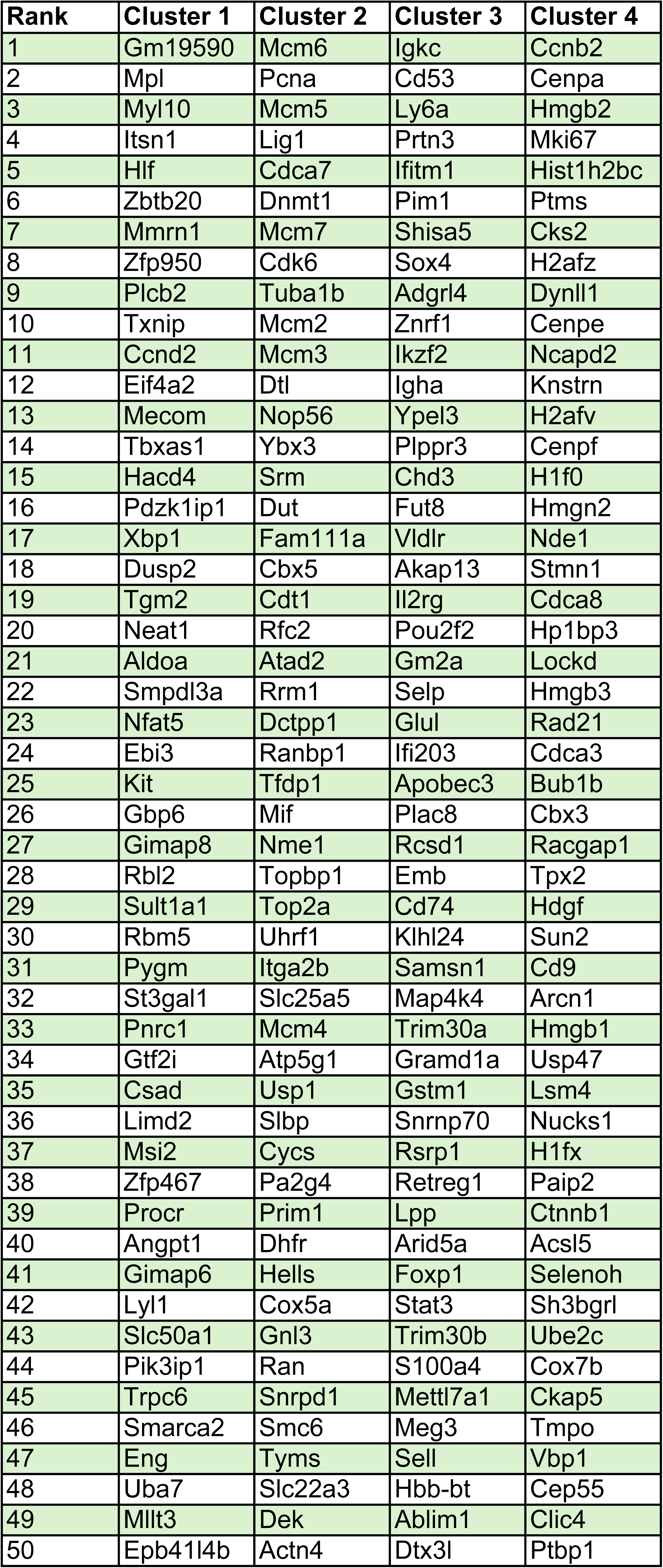
List of top50 genes differentially expressed in the 4 identified clusters.

**Table S2.**
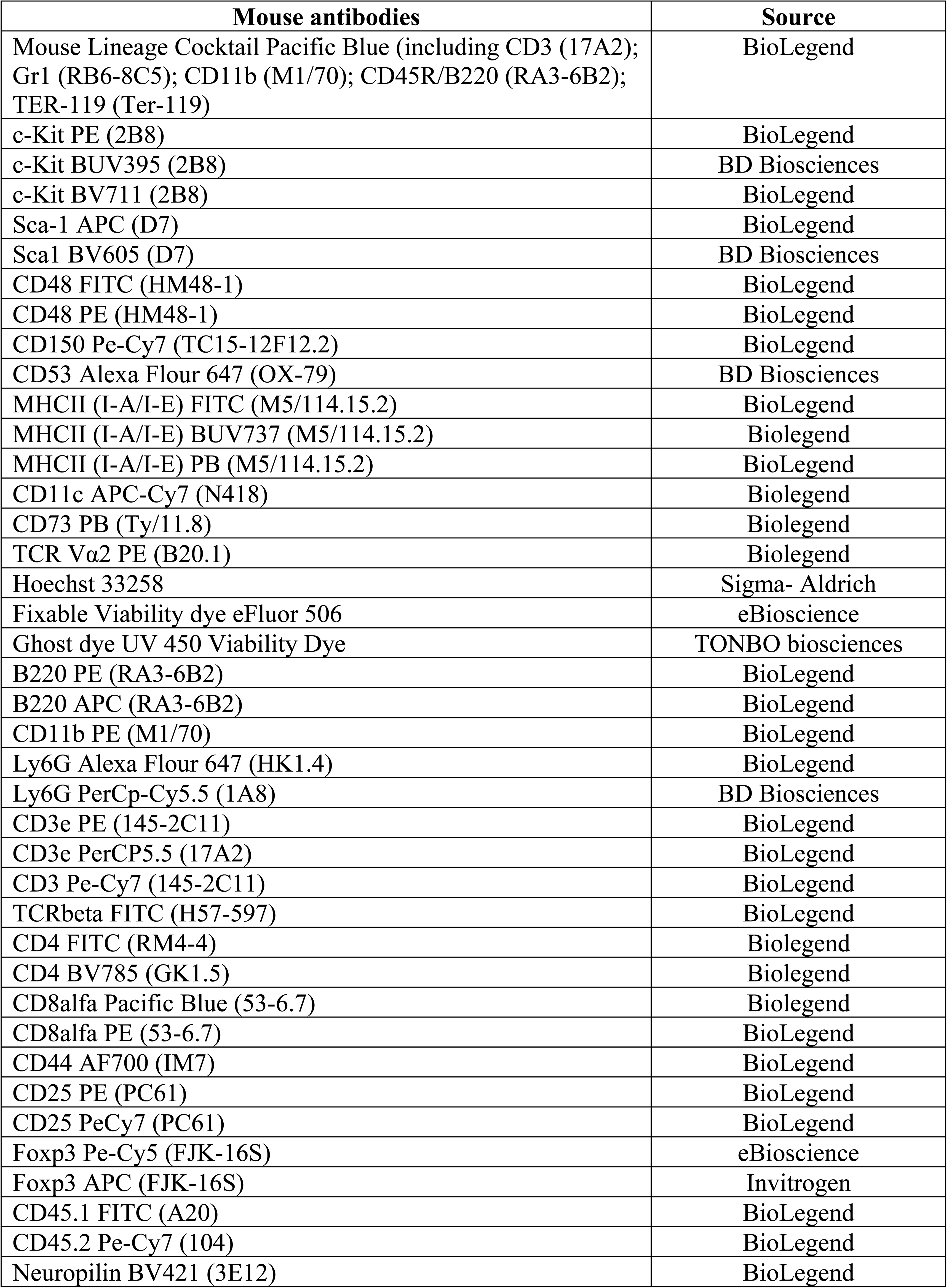
List of flow cytometry antibodies used.

**(*1*)**

