## Supplemental figures and Tables for "HSCs and Tregs cooperate to preserve extramedullary hematopoiesis under chronic inflammation"

Figure S1

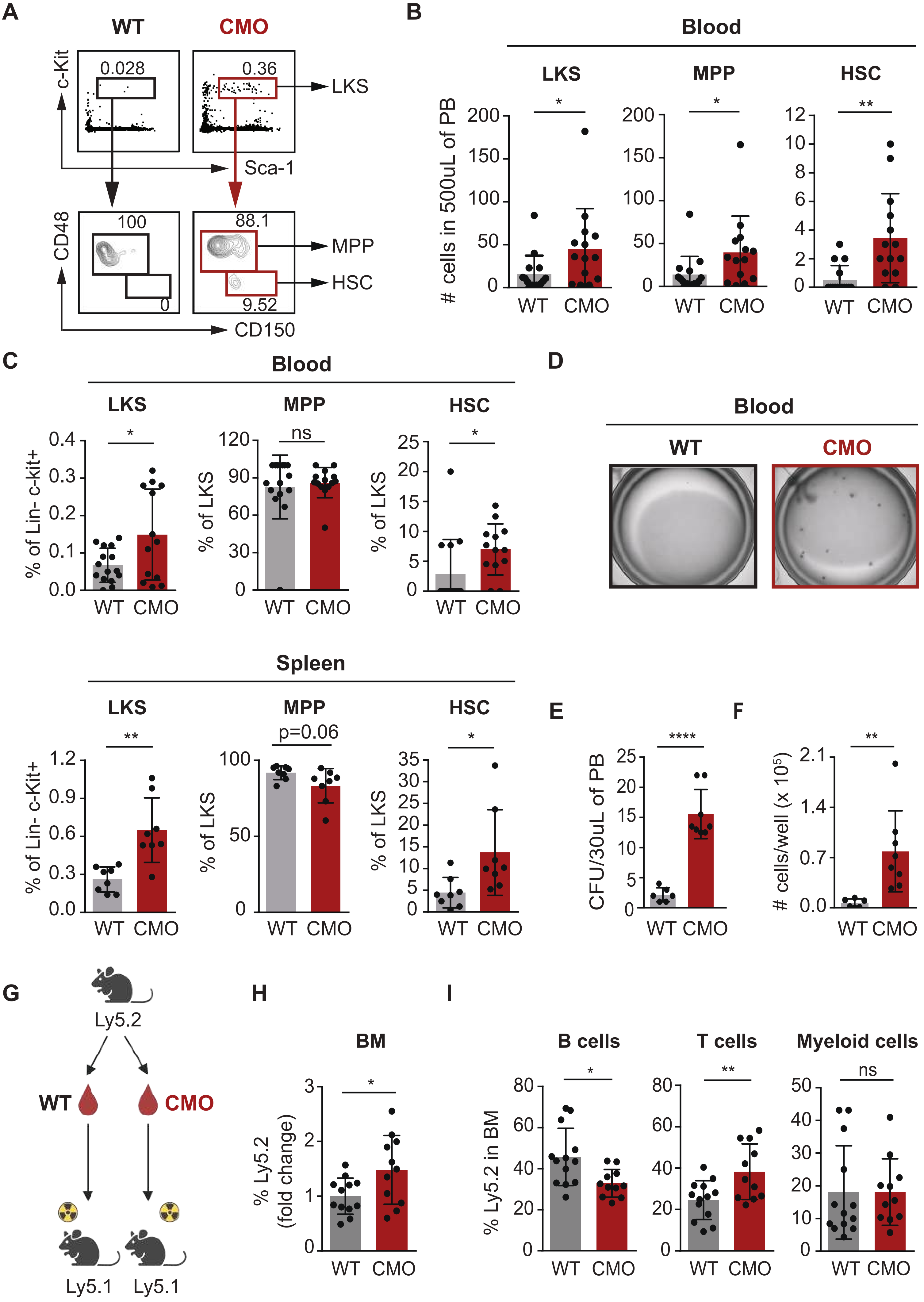

A
WT
CMO
c-Kit
0.028
0.36
LKS
Sca-1
CD48
100
0
88.1
9.52
MPP
HSC
CD150
B
Blood
LKS
MPP
HSC
### cells in 500uL of PB
*
*
**
WT
CMO
C
Blood
LKS
MPP
HSC
% of Lin- c-kit+
% of LKS
ns
*
D
Blood
WT
CMO
Spleen
LKS
MPP
HSC
% of Lin- c-Kit+
% of LKS
**
p=0.06
*
E
CFU/30uL of PB
****
F
### cells/well (x 10⁵)
**
G
Ly5.2
WT
CMO
Ly5.1
Ly5.1
H
BM
% Ly5.2 (fold change)
*
I
B cells
T cells
Myeloid cells
% Ly5.2 in BM
*
**
ns

**Supplementary Figure 1. CMO mice exhibit increased numbers of functional HSPCs in peripheral blood.**

**(A)** Representative flow cytometry plots from blood isolated from 1 WT and 1 CMO mouse. Upper plots illustrate c-Kit and Sca-1 expression in lineage- (Lin-) cells and the rectangle gates for Lin- c-Kit+ Sca-1+ (LKS) cells. Lower plots indicate CD48 and CD150 expression in LKS, and boxes illustrate gating for LKS CD48+ CD150- cells (MPPs) and LKS CD48- CD150+ cells (HSCs). Numbers indicate percentage from parental gates. **(B)** Quantification of panel a. Y-axes indicate the number of LKS, MPP, and HSC in 500 µL of peripheral blood (PB) from WT (gray) and CMO (red) mice. At least 8 mice were included per group, and each mouse is represented by a dot symbol. **(C)** Frequency of LKS, MPP, and HSC in blood and SP from WT (gray) and CMO (red) mice. Y-axes indicate percentage (%) from parental gate. Each dot indicates values for 1 mouse. All animals included were 16 to 20 weeks old. Data indicate mean ± SD from at least 2 independent experiments, and 2-tailes Student t test was used to assess statistical significance (*P , 0.05, **P , 0.01). **(D)** Representative microscopy images of colony culture assays using MethoCult M3434. A total of 30 µL of PB from WT (gray) and CMO (red) mice were plated per well. Images correspond to day 7 of culture. **(E-F)** Number of colony-forming units (CFU) (E) and cells (F) enumerated in panel c. Y-axes indicate the numbers per well at day 7. PB from at least 6 mice in 3 independent experiments was used in each condition. Each mouse is represented by a dot symbol. **(G)** Schematic representation of blood trans-plantation assays. Cells present in 1000 µL of PB from WT or CMO mice (Ly5.2) were transplanted into lethally irradiated Ly5.1 recipient mouse along with 0.5x106 BM support cells (Ly5.1). **(H)** Quantification of engraftment 16 weeks post-transplantation. Y-axes indicate percentage of WT (gray) and CMO (red) donor-derived Ly5.2+ cells in PB and BM. Engraftment is indicated as fold change from WT group. At least 11 animals were included in each group. Each dot indicates values for 1 animal. **(I)** Lineage reconstitution analysis 16 weeks after transplantation in BM from recipients transplanted with blood from WT (gray) and CMO (red) mice. Y-axis indicates the percentage of donor-derived Ly5.2+ B cells, T cells, and myeloid cells. Each dot indicates values for 1 recipient mouse. At least 11 recipients were used in each group. All animals included in Figure 1 were 12 to 25 weeks old. Data indicate mean ± SD from at least 3 indepen-dent experiments, and 2-tailed Student t test was used to assess statistical significance (*P , 0.05, **P , 0.01, ****P , 0.0001, and ns, not significant)

#### Figure S2

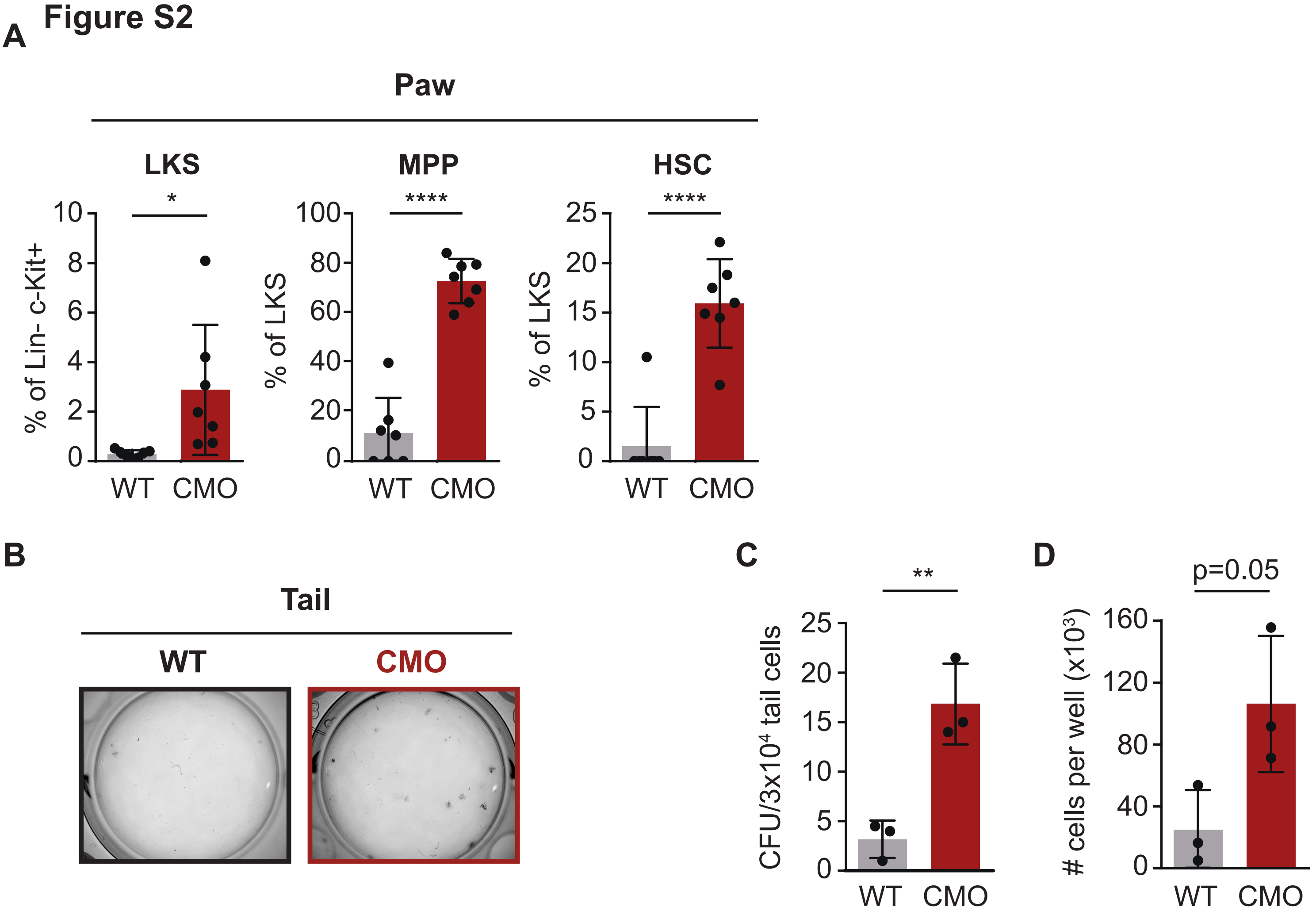


**Supplementary Figure 2. Presence of functional HSPCs in inflamed paws and tails.**

**(A)** Frequency of LKS (Lin- c-Kit+ Sca1+), MPP (Lin- c-Kit+ Sca1+ CD48+CD150-) and HSC (Lin- c-Kit + Sca1+ CD48-CD 150+) from paw. Y-axis indicates percentage (%) from parental gate in paw from WT (gray) and CMO (red) mice. Each dot indicates values for 1 mouse. All animals included were 16 to 20 weeks old. Data indicate mean ± SD from at least 3 independent experiments, and 2-tailed Student t test was used to assess statistical significance (*P , 0.05, ****P , 0.0001). **(B)** Representative microscopy images of colony culture assays after 7 days of culture. 3x103 WT (gray) and CMO (red) cells from tail were plated per well using MethoCult M3434. **(C)** Enumeration of panel b. Number of colony forming units (CFU) from cells isolated from WT (gray) and CMO (red) tails. Each dot symbol indicates values for one mouse. 2-tailed Student t test was used to assess statistical significance (**P , 0.01). **(D)** Number of cells in cultures from panel b. Y-axes indicate numbers of cells per well at day 7. Tail cell suspensions from 3 mice were used in each condition.

**Figure S3**

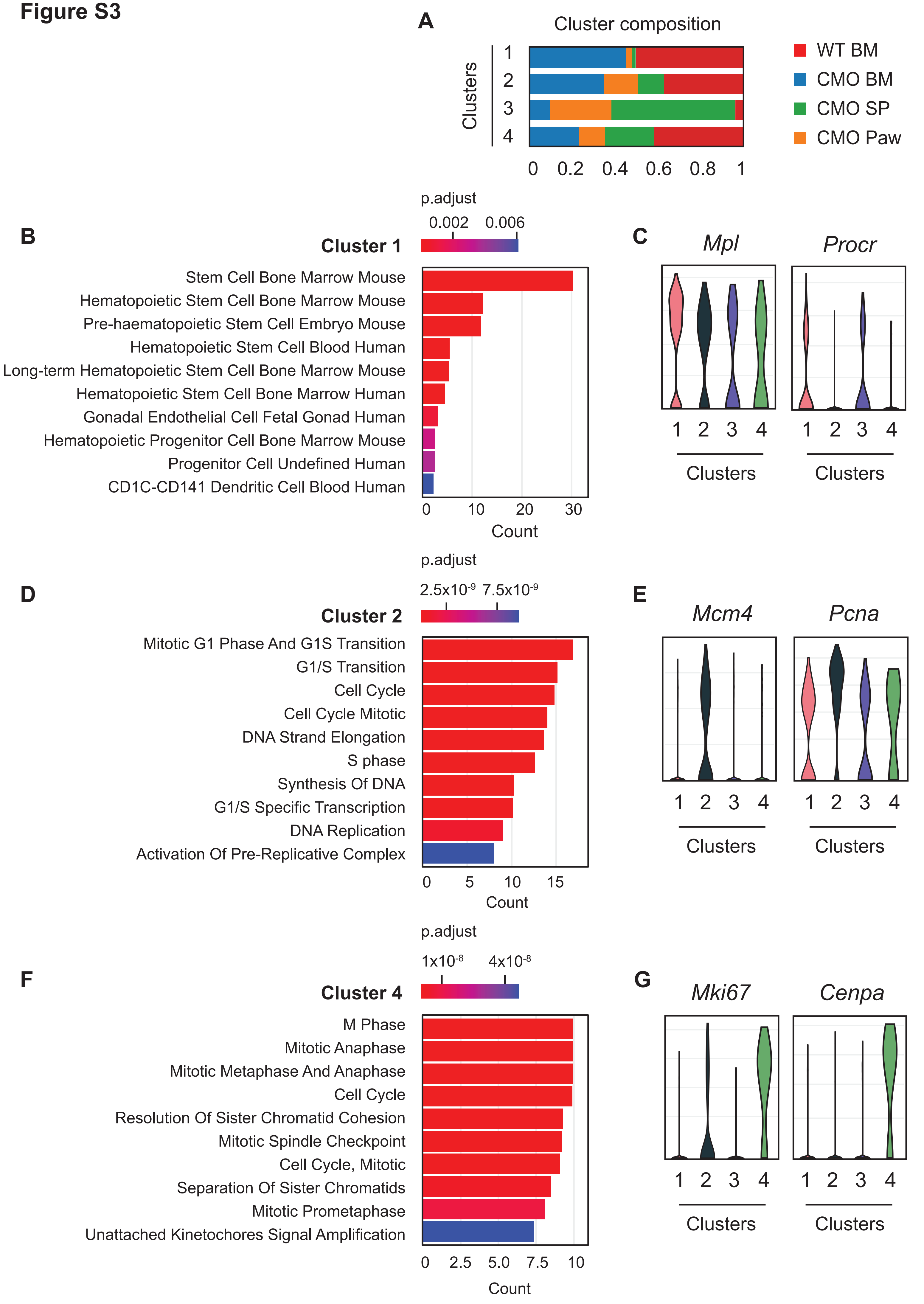
A
Cluster composition
Clusters
1
2
3
4
0
0.2
0.4
0.6
0.8
1
WT BM
CMO BM
CMO SP
CMO Paw
B
Cluster 1
p.adjust
0.002
0.006
Stem Cell Bone Marrow Mouse
Hematopoietic Stem Cell Bone Marrow Mouse
Pre-haematopoietic Stem Cell Embryo Mouse
Hematopoietic Stem Cell Blood Human
Long-term Hematopoietic Stem Cell Bone Marrow Mouse
Hematopoietic Stem Cell Bone Marrow Human
Gonadal Endothelial Cell Fetal Gonad Human
Hematopoietic Progenitor Cell Bone Marrow Mouse
Progenitor Cell Undefined Human
CD1C-CD141 Dendritic Cell Blood Human
0
10
20
30
Count
C
Mpl
Procr
1 2 3 4
Clusters
D
Cluster 2
p.adjust
2.5x10-9
7.5x10-9
Mitotic G1 Phase And G1S Transition
G1/S Transition
Cell Cycle
Cell Cycle Mitotic
DNA Strand Elongation
S phase
Synthesis Of DNA
G1/S Specific Transcription
DNA Replication
Activation Of Pre-Replicative Complex
0
5
10
15
Count
E
Mcm4
Pcna
1 2 3 4
Clusters
F
Cluster 4
p.adjust
1x10-8
4x10-8
M Phase
Mitotic Anaphase
Mitotic Metaphase And Anaphase
Cell Cycle
Resolution Of Sister Chromatid Cohesion
Mitotic Spindle Checkpoint
Cell Cycle, Mitotic
Separation Of Sister Chromatids
Mitotic Prometaphase
Unattached Kinetochores Signal Amplification
0
2.5
5.0
7.5
10
Count
G
Mki67
Cenpa
1 2 3 4
Clusters

**Supplementary Figure 3. Identification and classification of distinct HSCs clusters.**

**(A)** Relative contribution of WT BM, CMO BM, CMO SP and CMO paw HSCs to the cellular composition of each cluster. Y-axis indicates the cluster number. X-axis indicates the relative contribution of the individual HSC samples to each cluster.
**(B)** Relevant results of enrichment analysis of differentially expressed genes by Enrichr 1 in cluster 1.
**(C)** Violin plots representing the expression of Mpl and Procr genes in cluster 1.
**(D)** Relevant results of enrichment analysis of differentially expressed genes by Enrichr 1 in cluster 2.
**(E)** Violin plots representing the expression of Mcm4 and Pcna genes in cluster 2.
**(F)** Relevant results of enrichment analysis of differentially expressed genes by Enrichr 1 in cluster 4.
**(G)** Violin plots representing the expression of Mki67 and Cenpa genes in cluster 4.

#### Figure S4

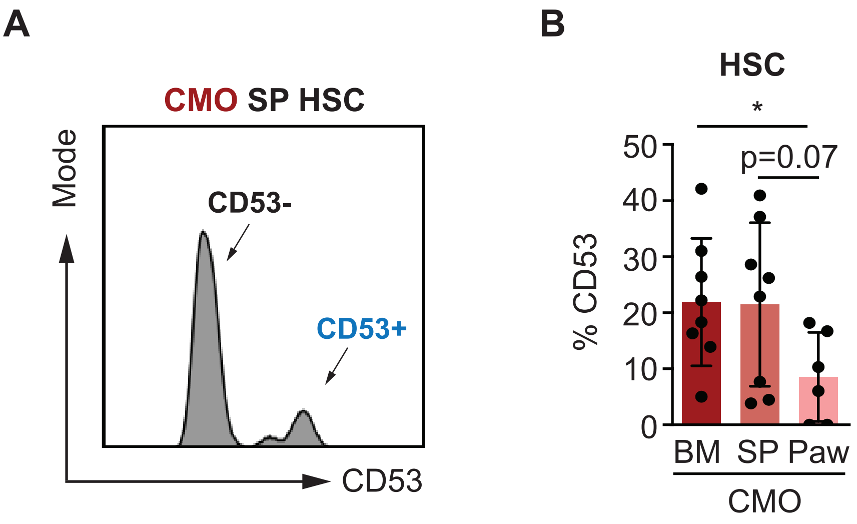


**Supplementary Figure 4. CD53 and Ki-67 labeling in HSCs.**

**(A)** Representative histogram plot of surface CD53 in CMO SP HSCs. Arrows indicate a subset of CD53- and a CD53+ HSCs.
**(B)** Frequency of CD53 in BM, SP and paw HSCs from CMO mice. Y-axis indicates percentage (%) of CD53 in the cell surface of HSCs.
All animals included were 12 to 15 weeks old. Data indicate mean ± SD from 3 independent experiments. 2-tailed Student t test was used to assess statistical significance (*P , 0.05, **P , 0.01, and ns, not significant).

#### Figure S5

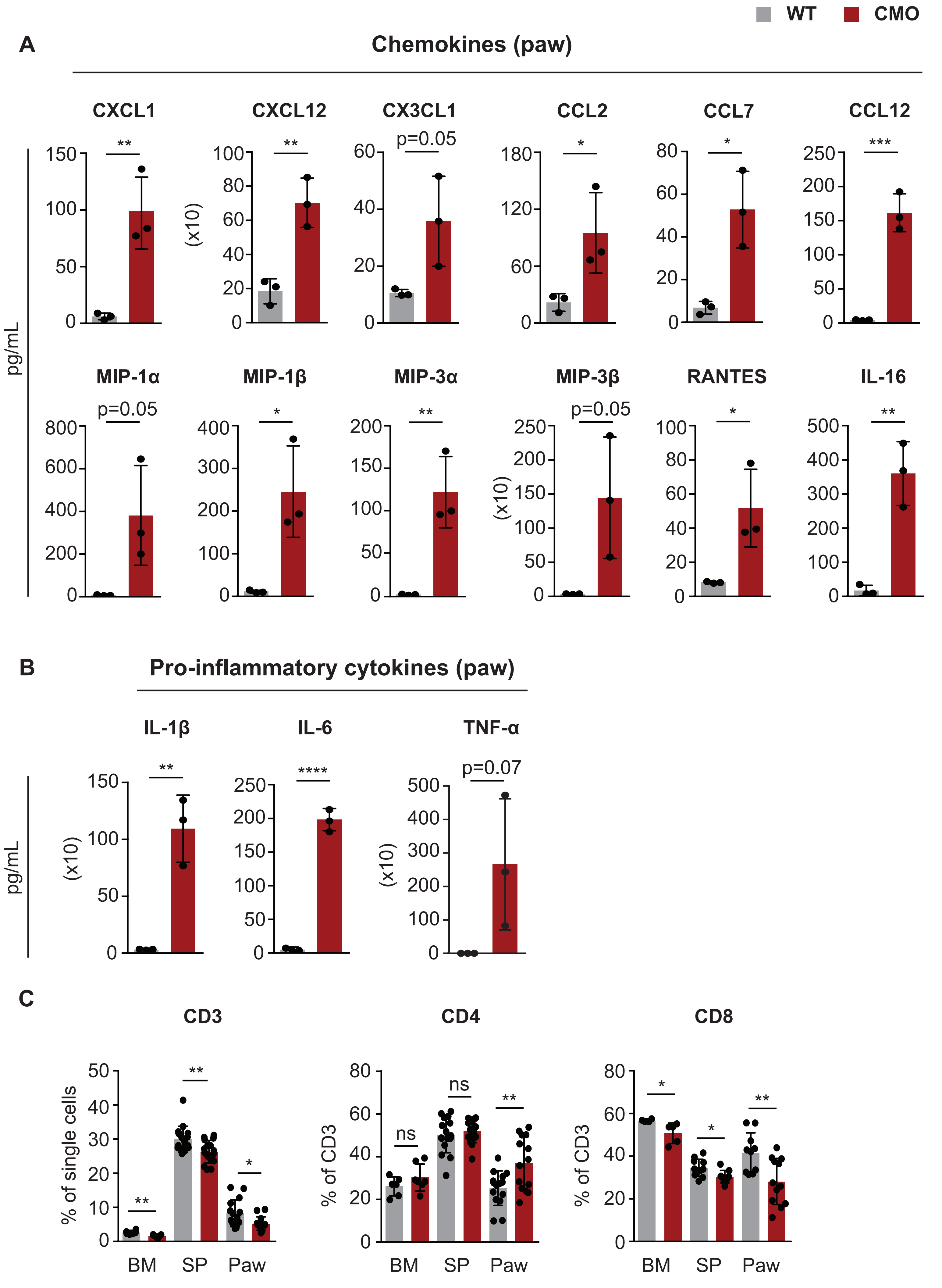

WT
CMO
A
Chemokines (paw)
CXCL1
CXCL12
CX3CL1
CCL2
CCL7
CCL12
pg/mL
(x10)
p=0.05
MIP-1α
MIP-1β
MIP-3α
MIP-3β
RANTES
IL-16
p=0.05
p=0.05
B
Pro-inflammatory cytokines (paw)
IL-1β
IL-6
TNF-α
p=0.07
pg/mL
(x10)
C
CD3
CD4
CD8
% of single cells
% of CD3
BM
SP
Paw
ns

**Supplementary Figure 5. Pro- and anti-inflammatory environment in CMO paws.**

**(A-B)** Chemokine and cytokine profiling by BioPlex in CMO paw. Each dot symbol indicates values for 1 mouse. Data indicate mean ± SD from 1 independent experiments.
**(C)** Frequency of CD3, CD4 and CD8 in BM, SP and paw HSCs from CMO mice. Y-axis indicates percentage (%) of CD3, CD4 or CD8 from parental gate. Each dot symbol indicates values for 1 mouse. Data indicate mean ± SD from 2 and more independent experiments.
All animals included were 16 to 25 weeks old. 2-tailed Student t test was used to assess statistical significance (*P , 0.05, **P , 0.01, ****P , 0.0001).

#### Figure S6

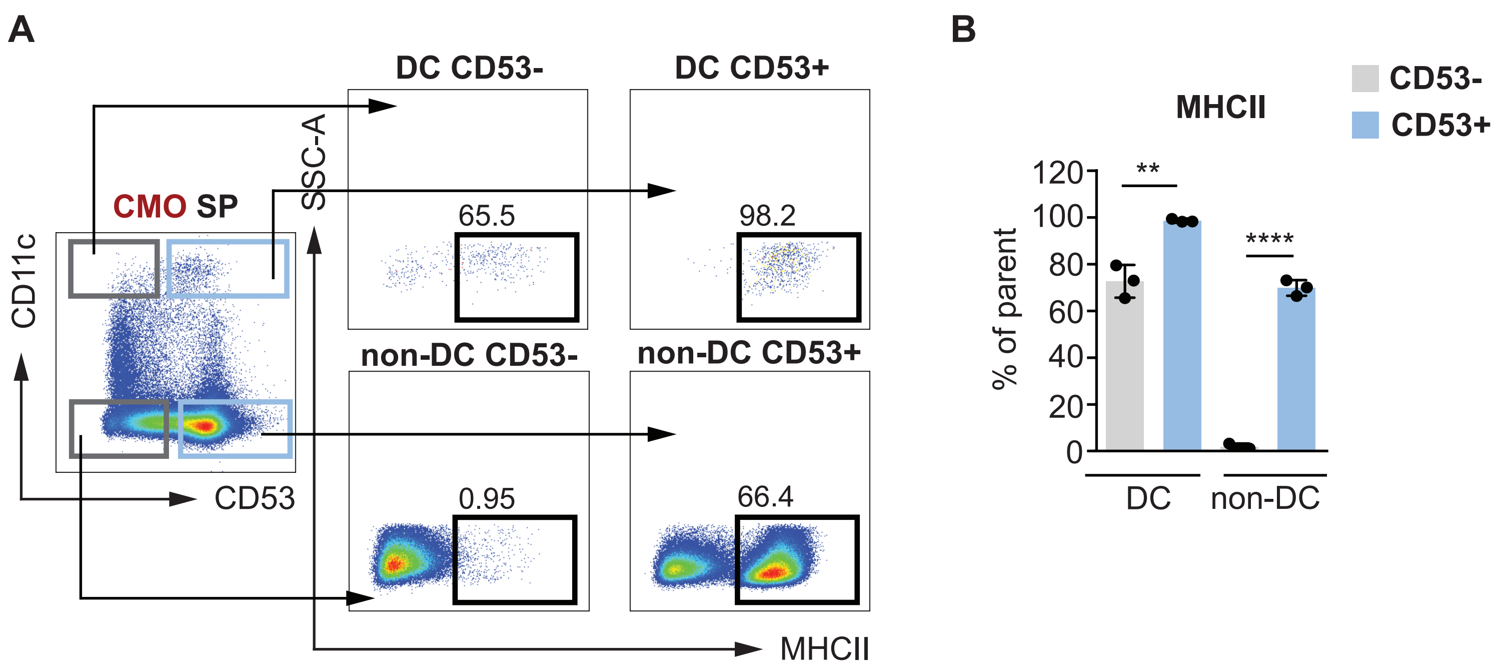


**Supplementary Figure 6. CD53+ cells contain higher levels of MHCII.**

**(A)** Representative gating strategy for MHCII in CMO SP. Splenocytes were gated according to CD11c and CD53 levels. Upper left box indi cates CD53- dentritic cells (DC CD53-), upper right box CD53+ CD11c+ cells (DC CD53+), lower left box CD53- CD11c- (non-DC CD53-), and lower right box CD53+ CD11c- (non-DC CD53+). The rest of the dot plots indicate MHCII levels in the distinct populations. Black boxes indicate MHCII+ cells. Numbers indicate percentages from parental gates. (B) Frequency of MHCI lexpression across in DC and non-DC pop ulations from CMO SP. CD53- cells are indicated in gray and CD53+ in blue.

Each dot symbol indicates values for 1 mouse. Data indicate mean ± SD from 1 experiment. 2-tailed Student t test was used to assess statistical significance (**P , 0.01, ****P , 0.0001).

**Figure S7**

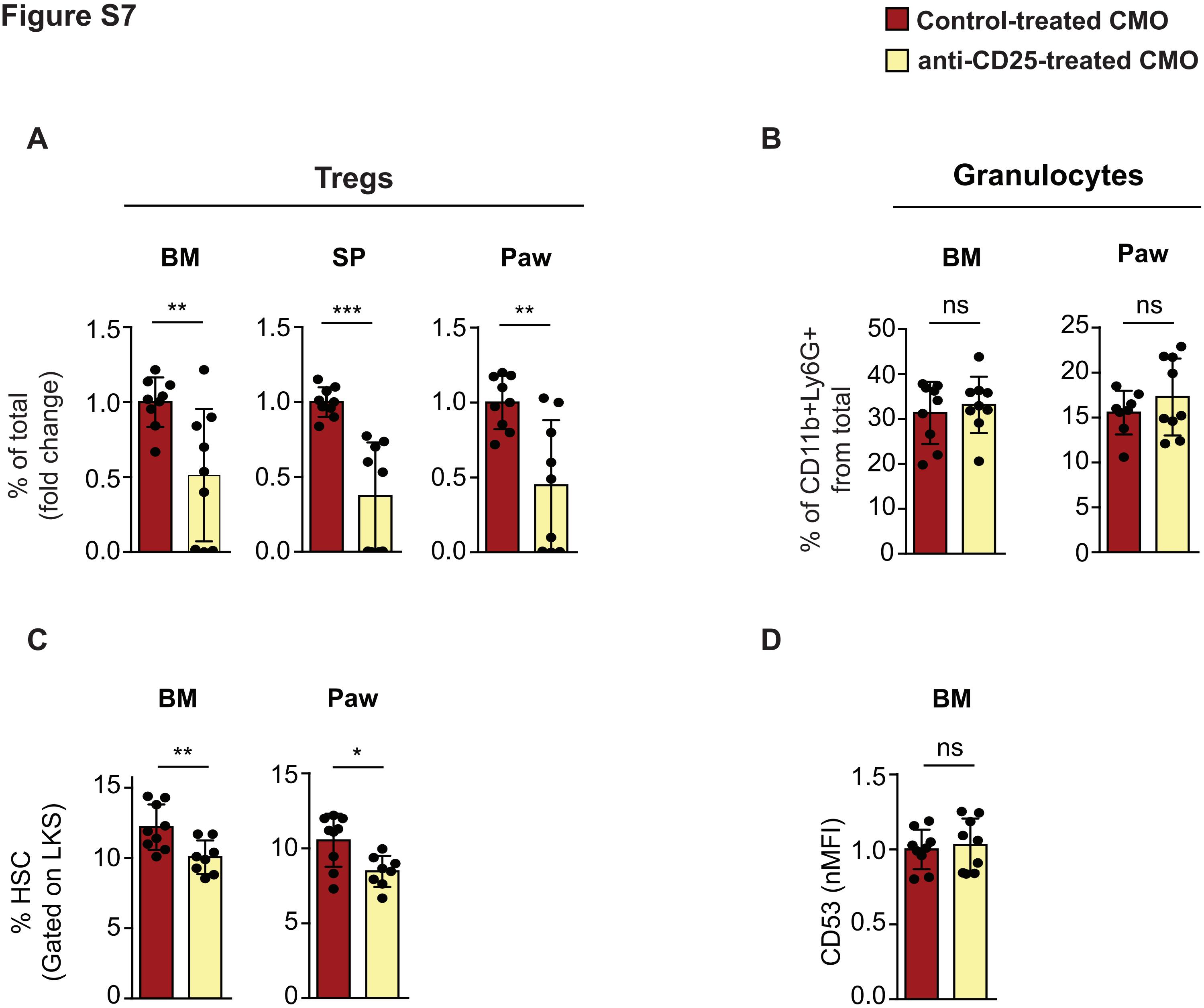


**Supplementary Figure 7. Treg depletion effect on BM and paw HSC.**

**(A)** Frequency of CD25+Foxp3+ Tregs in non-treated (red) and anti-CD25 treated (yellow) CMO mice in BM, SP and paw. Y-axis indicates percentage (%) of Tregs from total cells and is calculated as a fold change from non-treated (red) CMO group. X-axis indicates whether mice received (yellow) or not (red) anti-CD25 antibody treatment.
**(B)** Frequency of CD11b+ Ly6G+ granulocytes in non-treated (red) and anti-CD25 treated (yellow) CMO mice in BM and paw. Y-axis indicates percentage (%) of granulocytes from parental gate. X-axis indicates whether mice received (yellow) or not (red) anti-CD25 antibody treatment.
**(C)** Frequency of HSC (Lin- c-Kit+ Sca1+ CD48-CD 150+) from control (red) or anti-CD25 (yellow) CMO treated mice in BM and paw. Y-axis indicates percentage (%) from parental LKS gate. X-axis indicates whether mice received (yellow) or not (red) anti-CD25 antibody treatment.
**(D)** Flow cytometric analysis of total CD53 expression in HSC from CMO BM. HSCs were isolated from non-treated (red) and anti-CD25-treated (yellow) CMO mice. Total CD53 expression is indicated as a fold change from non-treated (red) CMO group. Y-axis indicates CD53 mean fluorescence intensity (MFI). X-axis indicates whether mice received (yellow) or not (red) anti-CD25 antibody treatment. Each dot indicates values for 1 mouse. Data indicate mean ± SD from 3 independent experiments. 2-tailed Student t test was used to assess statistical significance (ns, not significant).

| Rank | Cluster 1 | Cluster 2 | Cluster 3 | Cluster 4 |
|---|---|---|---|---|
| 1 | Gm19590 | Mcm6 | Igkc | Ccnb2 |
| 2 | Mpl | Pcna | Cd53 | Cenpa |
| 3 | Myl10 | Mcm5 | Ly6a | Hmgb2 |
| 4 | Itsn1 | Lig1 | Prtn3 | Mki67 |
| 5 | Hlf | Cdca7 | Ifitm1 | Hist1h2bc |
| 6 | Zbtb20 | Dnmt1 | Pim1 | Ptms |
| 7 | Mmrn1 | Mcm7 | Shisa5 | Cks2 |
| 8 | Zfp950 | Cdk6 | Sox4 | H2afz |
| 9 | Plcb2 | Tuba1b | Adgrl4 | Dynll1 |
| 10 | Txnip | Mcm2 | Znrf1 | Cenpe |
| 11 | Ccnd2 | Mcm3 | Ikzf2 | Ncapd2 |
| 12 | Eif4a2 | Dtl | Igha | Knstrn |
| 13 | Mecom | Nop56 | Ypel3 | H2afv |
| 14 | Tbxas1 | Ybx3 | Plppr3 | Cenpf |
| 15 | Hacd4 | Srm | Chd3 | H1f0 |
| 16 | Pdzk1ip1 | Dut | Fut8 | Hmgn2 |
| 17 | Xbp1 | Fam111a | Vldlr | Nde1 |
| 18 | Dusp2 | Cbx5 | Akap13 | Stmn1 |
| 19 | Tgm2 | Cdt1 | Il2rg | Cdca8 |
| 20 | Neat1 | Rfc2 | Pou2f2 | Hp1bp3 |
| 21 | Aldoa | Atad2 | Gm2a | Lockd |
| 22 | Smpdl3a | Rrm1 | Selp | Hmgb3 |
| 23 | Nfat5 | Dctpp1 | Glul | Rad21 |
| 24 | Ebi3 | Ranbp1 | Ifi203 | Cdca3 |
| 25 | Kit | Tfdp1 | Apobec3 | Bub1b |
| 26 | Gbp6 | Mif | Plac8 | Cbx3 |
| 27 | Gimap8 | Nme1 | Rcsd1 | Racgap1 |
| 28 | Rbl2 | Topbp1 | Emb | Tpx2 |
| 29 | Sult1a1 | Top2a | Cd74 | Hdgf |
| 30 | Rbm5 | Uhrf1 | Klhl24 | Sun2 |
| 31 | Pygm | Itga2b | Samsn1 | Cd9 |
| 32 | St3gal1 | Slc25a5 | Map4k4 | Arcn1 |
| 33 | Pnrc1 | Mcm4 | Trim30a | Hmgb1 |
| 34 | Gtf2i | Atp5g1 | Gramd1a | Usp47 |
| 35 | Csad | Usp1 | Gstm1 | Lsm4 |
| 36 | Limd2 | Slbp | Snrnp70 | Nucks1 |
| 37 | Msi2 | Cycs | Rsrp1 | H1fx |
| 38 | Zfp467 | Pa2g4 | Retreg1 | Paip2 |
| 39 | Procr | Prim1 | Lpp | Ctnnb1 |
| 40 | Angpt1 | Dhfr | Arid5a | Acsl5 |
| 41 | Gimap6 | Hells | Foxp1 | Selenoh |
| 42 | Lyl1 | Cox5a | Stat3 | Sh3bgrl |
| 43 | Slc50a1 | Gnl3 | Trim30b | Ube2c |
| 44 | Pik3ip1 | Ran | S100a4 | Cox7b |
| 45 | Trpc6 | Snrpd1 | Mettl7a1 | Ckap5 |
| 46 | Smarca2 | Smc6 | Meg3 | Tmpo |
| 47 | Eng | Tyms | Sell | Vbp1 |
| 48 | Uba7 | Slc22a3 | Hbb-bt | Cep55 |
| 49 | Mllt3 | Dek | Ablim1 | Clic4 |
| 50 | Epb41l4b | Actn4 | Dtx3l | Ptbp1 |

**Table S1.** List of top50 genes differentially expressed  in the 4 identified clusters

| Mouse antibodies | Source |
|---|---|
| Mouse Lineage Cocktail Pacific Blue (including CD3 (17A2); Gr1 (RB6-8C5); CD11b (M1/70); CD45R/B220 (RA3-6B2); TER-119 (Ter-119) | BioLegend |
| c-Kit PE (2B8) | BioLegend |
| c-Kit BUV395 (2B8) | BD Biosciences |
| c-Kit BV711 (2B8) | BioLegend |
| Sca-1 APC (D7) | BioLegend |
| Sca1 BV605 (D7) | BD Biosciences |
| CD48 FITC (HM48-1) | BioLegend |
| CD48 PE (HM48-1) | BioLegend |
| CD150 Pe-Cy7 (TC15-12F12.2) | BioLegend |
| CD53 Alexa Flour 647 (OX-79) | BD Biosciences |
| MHCII (I-A/I-E) FITC (M5/114.15.2) | BioLegend |
| MHCII (I-A/I-E) BUV737 (M5/114.15.2) | Biolegend |
| MHCII (I-A/I-E) PB (M5/114.15.2) | BioLegend |
| CD11c APC-Cy7 (N418) | Biolegend |
| CD73 PB (Ty/11.8) | Biolegend |
| TCR Vα2 PE (B20.1) | Biolegend |
| Hoechst 33258 | Sigma- Aldrich |
| Fixable Viability dye eFluor 506 | eBioscience |
| Ghost dye UV 450 Viability Dye | TONBO biosciences |
| B220 PE (RA3-6B2) | BioLegend |
| B220 APC (RA3-6B2) | BioLegend |
| CD11b PE (M1/70) | BioLegend |
| Ly6G Alexa Flour 647 (HK1.4) | BioLegend |
| Ly6G PerCp-Cy5.5 (1A8) | BD Biosciences |
| CD3e PE (145-2C11) | BioLegend |
| CD3e PerCP5.5 (17A2) | BioLegend |
| CD3 Pe-Cy7 (145-2C11) | BioLegend |
| TCRbeta FITC (H57-597) | BioLegend |
| CD4 FITC (RM4-4) | Biolegend |
| CD4 BV785 (GK1.5) | Biolegend |
| CD8alfa Pacific Blue (53-6.7) | Biolegend |
| CD8alfa PE (53-6.7) | BioLegend |
| CD44 AF700 (IM7) | BioLegend |
| CD25 PE (PC61) | BioLegend |
| CD25 PeCy7 (PC61) | BioLegend |
| Foxp3 Pe-Cy5 (FJK-16S) | eBioscience |
| Foxp3 APC (FJK-16S) | Invitrogen |
| CD45.1 FITC (A20) | BioLegend |
| CD45.2 Pe-Cy7 (104) | BioLegend |
| Neuropilin BV421 (3E12) | BioLegend |

**Table S2. List of flow cytometry antibodies used.**
